# RB1 gene mutations and their association with Endometrial Cancer

**DOI:** 10.1101/2025.09.22.677485

**Authors:** Pritam Kumar Ghosh, Somrita Das, S N Gagan Gaurav, Sahana Ghosh, Arindam Maitra, Manjula Das, Saumitra Das

## Abstract

Retinoblastoma gene (RB1) mutation has been reported in lung cancer, retinoblastoma and cervical cancer etc. Here, we elucidate the involvement of RB1 mutations and their regulation in endometrial cancer. Analysis of mutation data of 547 endometrial cancer samples revealed that 12% of samples harboured RB1 mutations. However, 26% of the patients aged between 30 and 50 years harboured RB1 mutations, compared to only 10% in the older age group. Further, in silico interaction studies and structural analysis of these novel RB1 mutations and E2F revealed possible physiological relevance. We found 14,117 genes were significantly mutated, and 5,770 genes were differentially expressed in RB1-altered patients. Pathway analysis showed a significant correlation between RB1 mutations and estrogen receptor mutations. Different cancer-related pathways were also altered in RB1-mutated conditions. More importantly, a positive link between RB1 mutations and HPV infection pathways was observed, indicating HPV infections might induce RB1 mutations in endometrial cancer, which could be prevented by early vaccination. Our findings suggest RB1 mutations as a potential contributor to endometrial cancer in younger women, highlighting the importance of viral co-factors in risk assessments, advocating HPV vaccination in individuals with RB1 mutations.

## 1. Introduction

The Retinoblastoma gene (RB1), a well-studied tumour suppressor gene, is a negative regulator of the cell cycle and cancer. The RB1 gene is located on chromosome 13q14.2 and has a total length of 178143 bp, which contains 27 coding exons, 26 introns and a core promoter, encoding a 105-kD (928 amino acids) nuclear phosphoprotein, pRB (retinoblastoma protein) (1,2). pRB belongs to a larger gene family that includes two other genes, retinoblastoma-like 1 (RBL1/p107) and retinoblastoma-like 2 (RBL2/p130). It comprises 3 domains: RbN, RbAB (pocket domain), and RbC (3).

The biological functions of Rb1 include tumour suppression, differentiation, and apoptosis. It is shown that pRB can interact with more than 100 proteins in the cell (4). pRB switches between a hyperphosphorylated and relatively unphosphorylated state in a cell cycle-specific manner. It is underphosphorylated in G1, becomes heavily phosphorylated just before the G1 to S transition, remains phosphorylated in S, G2, and most of M, and reverts to an underphosphorylated state at or before the M-G1 transition. (5). Unphosphorylated pRB binds to E2F1 and inhibits E2F1-mediated transcription of Cyclin A, Cyclin E and cdc25, thereby inducing cell cycle arrest (6). Mutant pRB cannot interact with the E2Fs, and thus leads to cell proliferation (7).

RB1 mutation is a widely recognized carcinogenic mutation that established a paradigm for oncologic research because of its early discovery, its recessive nature, and its high frequency of occurrence (8). RB1 gene mutations occur in various forms throughout its coding exons and introns. The most (approximately 85%) pathogenic mutations are located in all 27 exons and one core promoter region of the RB1 gene (9). RB1 protein is inactivated by different viral infections like Human Papillomavirus (HPV), Adenovirus, Simian Virus 40. (10–12). RB1 mutations are well studied in retinoblastoma, lung cancer, breast cancer, and prostate cancer (13–16). The current study aimed to elucidate the presence of RB1 mutations in other cancers and their implications in cancer progression, particularly in endometrial cancer, as RB1 mutations are frequently observed in this disease. Endometrial cancer was caused by genetic, familial, and lifestyle factors, and 329 genes were known to be genetically altered (17). Structural analysis of the RB1 mutation, associated clinical correlation, and altered gene pathway analysis were conducted to understand the cause and consequences of the cancer.

## 2. Methods

### 2.1 Patient datasets

This study was conducted using five publicly available cancer datasets. (i) Retinoblastoma cancer cohort (Retinoblastoma (MSK, Cancers 2021). This includes 83 samples from 81 patients (18). (ii) MSK MetTropism (MSK, Cell 2021) cohort, which includes 25,775 samples per patient across 27 different cancer types, including endometrial cancer (S1A) (19). Most of the samples were taken from the white race people (78.6%, S1B) in this dataset. Additionally, the endometrial cancer samples (n=1,315) from this cohort were analysed exclusively to determine its distinct genomic profile. (iii) Further, the endometrial cancer dataset (Endometrial Carcinoma-TCGA, GDC) was considered, which consisted of 548 samples from 547 patients, of which 68.4% of the samples were taken from white people (S1C). Samples from other gynaecological cancers were analysed too which were (iv) Cervical Squamous Cell Carcinoma (TCGA, PanCancer Atlas) and (v) High-Grade Serous Ovarian Cancer (TCGA, GDC). The Cervical Squamous Cell Carcinoma dataset contained 297 samples per patient, whereas the Ovarian Cancer dataset contained 602 samples from 586 patients. Both of the datasets also majorly contained white race people (S1E and S1D). For all the datasets, vivid genomic and clinical data were available, which were utilised for this study. Additionally, the transcriptomics data available for the cancer datasets under the TCGA study were analysed too. The data from the mentioned datasets are available in cbioportal (https://www.cbioportal.org/)

### 2.2 mRNA and Protein Expression analysis

The expression of RB1 mRNA and Protein in different tissues were checked in the human protein atlas(https://www.proteinatlas.org/). The expression of mRNA and Protein in cancer tissues was plotted using UALCAN (https://ualcan.path.uab.edu/). mRNA expression analysis of RB1 altered vs unaltered was done in cBioportal. UALCAN and cBioPortal utilize data from TCGA for mRNA expression analysis and from CPTAC for protein expression analysis, and provide statistical significance for their results. Molbiotools were used to compare different datasets (https://molbiotools.com/listcompare.php).

### 2.3 Protein Structure Preparation

The Protein Data Bank provided the crystallographic structure of the human retinoblastoma protein (Rb) in complex with the E2F1 transactivation domain (TAD), which was resolved at 2.20 Å and has PDB ID 1N4M. The vast pocket domain necessary for E2F binding, represented by Chain A of Rb (residues 373–928), was isolated and utilized as the wild-type (WT) reference for docking studies. Four Rb variants: E137* (premature stop codon), R272I, R741C, and R876C were modelled using the AlphaFold2-based ColabFold implementation (v1.5.2) with default parameters(20), no structural templates, and AMBER relaxation enabled in order to examine the impact of clinically significant mutations linked to endometrial cancer.

### 2.4 Modelling of E2F Proteins

Human E2F1 and E2F2 transactivation domains (TADs) were chosen for docking. AlphaFold2 was used to model residues that corresponded to the known Rb-binding area in the E2F family (E2F1: residues 409–437, E2F2: residues 410–438). The template_mode was set to none, the MSA pairing method was set to complete, and the msa_mode was set to MMseqs2 (unpaired). Steric conflicts were reduced by using AMBER relaxation. The pLDDT score was used to rank the models, and the model with the highest rating for each protein was chosen.

### 2.5 Docking and Analysis

In addition to literature and inherent disorder predictions, the crystal structure of the Rb–E2F1 complex (PDB: 1N4M) was used to identify interface residues for docking. Residues 746–761, 813–825, and 865–877 on the Rb protein (chain A) were chosen because they match helices in the B-domain that are essential for E2F binding. Regions surrounding the conserved marked box and Rb-binding motif were selected for E2F1 (residues 409–437) and E2F2 (residues 410– 438). For HADDOCK2.4, these residues were identified as active, while nearby residues were automatically classified as passive(21,22). On the HADDOCK2.4 online server, protein-protein docking was performed using the Rb wild-type (PDB 1N4M) and mutant models (E137*, R272I, R741C, and R876C from AlphaFold), as well as E2F1 and E2F2 structures predicted by AlphaFold. One thousand rigid-body models (it0), two hundred semi-flexible refinements (it1), and two hundred water-refined models were used in the docking process. Using a minimum size of 4 and a 1.0 Å RMSD limit, clustering was carried out using the fraction of common contacts (FCC). The HADDOCK score was used to rank the top clusters, and PyMOL v2.5 and Ligplot+ were used to choose the lowest-energy model from the best cluster for downstream interaction and structural analysis.

### 2.6 Pathway analysis

Pathway enrichment analysis of the genomic and transcriptomic data was performed using ShinyGO 0.82 (https://bioinformatics.sdstate.edu/go/). The top pathways were shortened by FDR (false discovery rate) and enrichment score. Additionally, the tool mapped the associated genes to their respective chromosomal locations.

### 2.7 Statistical analysis

For statistical significance, p-values and q-values were calculated. The p-values were adjusted for multiple testing by applying the Benjamini-Hochberg correction method. Two-sided Fisher’s Exact test was performed to derive the statistical significance of the genomic alternations between altered and unaltered group, Student’s t-test was performed to identify differential mRNA expression between groups. The significance of the differential abundance of Mutation count and fraction Genome Altered between clinically different groups was calculated by performing the Wilcoxon Rank Sum test.

## 3. Results

### 3.1 RB1 Mutations and levels in different cancers

As 45% of retinoblastomas were reported to be caused by mutations in the RB1 gene (13), RB1 mutation was checked in the Retinoblastoma (MSK, Cancers 2021) data set. 75% of the samples contained RB1 mutations (Figure 1A, Data sheet 1). Nonsense mutation was the most common one with the C→T transition (Figure S2A and S2B). R320*, a nonsense mutation, was found in 5 patients. RB1 mutations were also analysed in other cancers with a dataset containing 25,775 patients/samples (Data sheet 2). There were 7% RB1 mutations among the different cancers (Figure 1B). Out of 27 different cancers, small-cell lung cancer had the highest number of RB1 alterations (mutation, structural variant, amplification, deep deletion, etc.); sarcomas and bladder cancers were followed by. The 8^th^ highest number of RB1 was found in endometrial cancer, which is a novel observation (Figure 1C).

**Figure 1:**
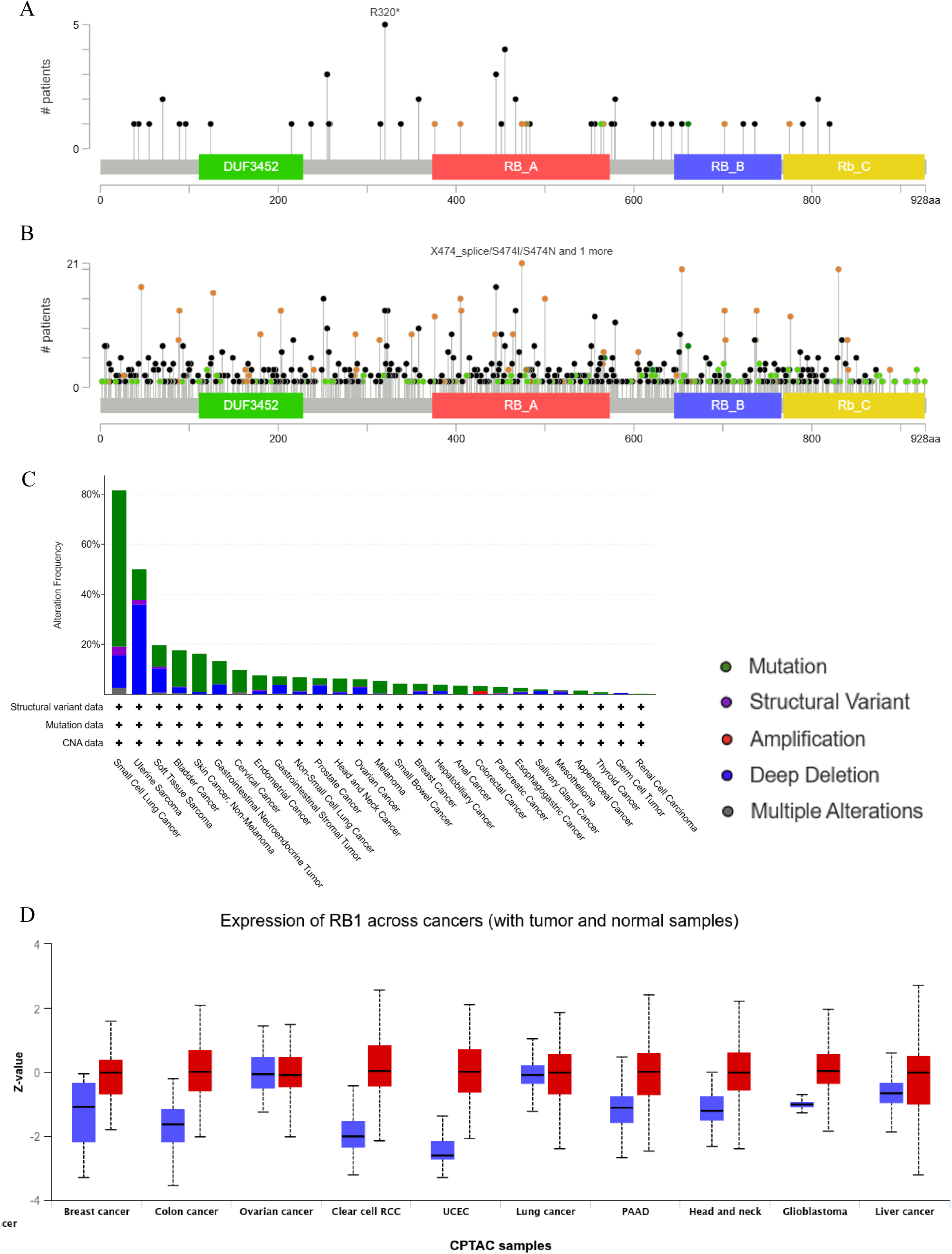
RB1 Mutations in different cancers. (A) Lollipop graph of RB1 mutations in retinoblastoma (cBioportal). (B) Lollipop graph of RB1 mutations in different cancers (cBioportal; MSK MetTropism dataset). (C) RB1 alterations in different cancers (MSK MetTropism dataset). (D) Expression of RB1 protein across the cancers (UALCAN).

Before understanding the relevance of RB1 mutations in different cancers, it is essential to understand the expression of RB1 in normal tissues and cancerous tissues. Other than the prostate, most of the tissues had high to medium expression of RB protein and RNA. The endometrium also contained a medium level of RB1 mRNA and protein (Figure S3A and S3B). The expressions of pRb were significantly upregulated in breast cancer, colon cancer, renal carcinoma, endometrial cancer, pancreatic cancer, head and neck cancer, glioblastoma and liver cancer (Figure 1D). Interestingly, pRB was mutated and significantly upregulated in endometrial cancer (fold change: p value = 9.09 × 10^-37^; Figure S3C). Therefore, in this study, we investigated a novel class of RB1 mutations in endometrial cancer, along with their possible causes and consequences.

### 3.2 RB1 mutations in endometrial cancer

RB1 mutations in endometrial cancer were analysed in 547 samples (Figure S1C, Data sheet 3), which contained genomics and transcriptomics data of all samples. The RB1 mutation was present in 12% of the samples (Figure 2A). The G→T transversion and Missense mutation were the most commonly found mutations in endometrial cancer (Figure 2B and 2C). Overall, the mutation count was significantly higher in the RB1-altered group (mutated group), indicating the presence of additional mutations in this group (Figure 2D). Patients in the lower age group (30-50 years) had higher frequencies (26%) of RB1 mutations compared to the older age group (10%) (Figure 2E; Data sheet 14). The E137* nonsense mutation was the most common mutation in the endometrial cancer that made a truncated protein (Figure 2F). Structural studies of the novel mutations were done by modelling and interaction of E2Fs was done by molecular docking.

**Figure 2:**
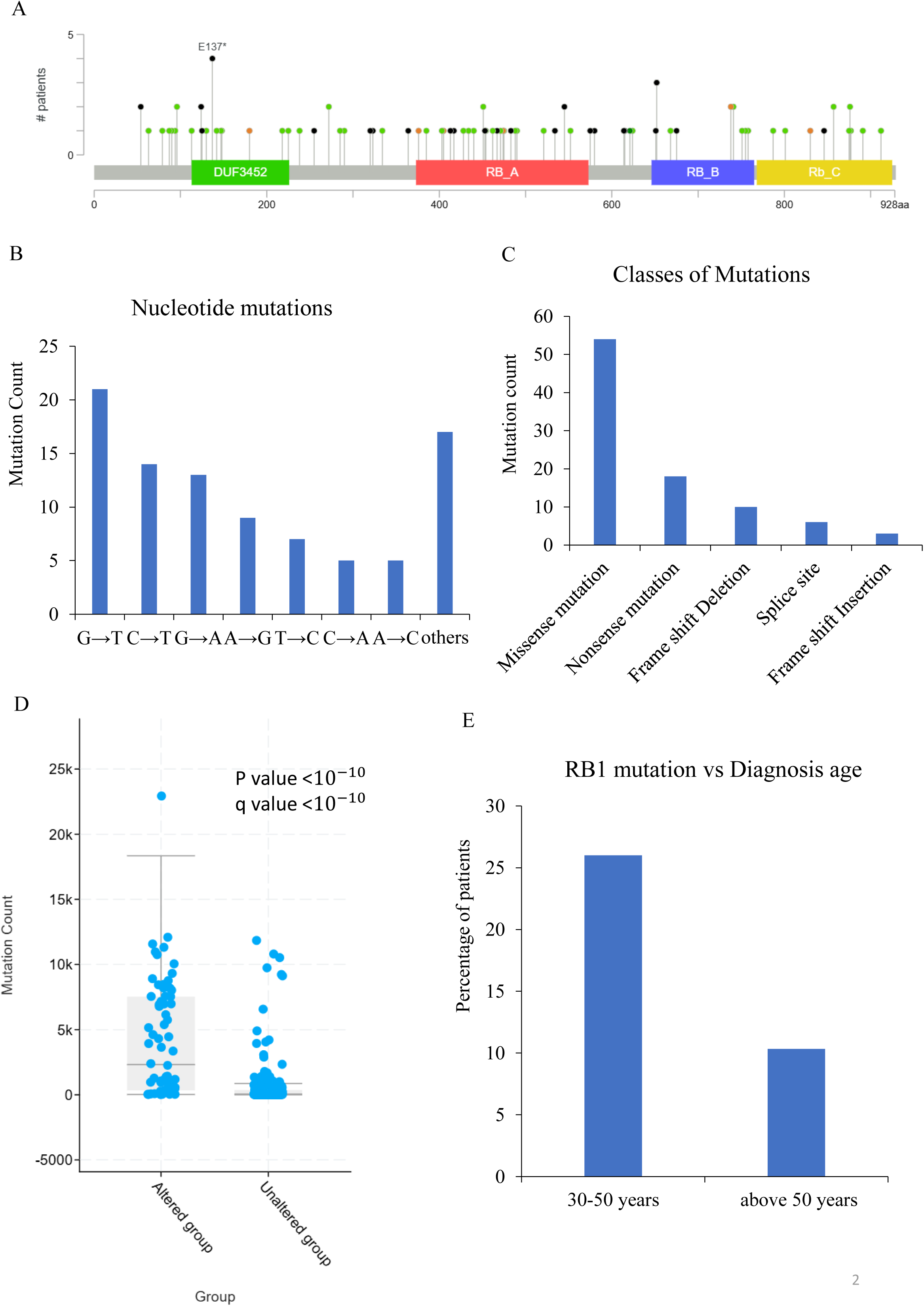
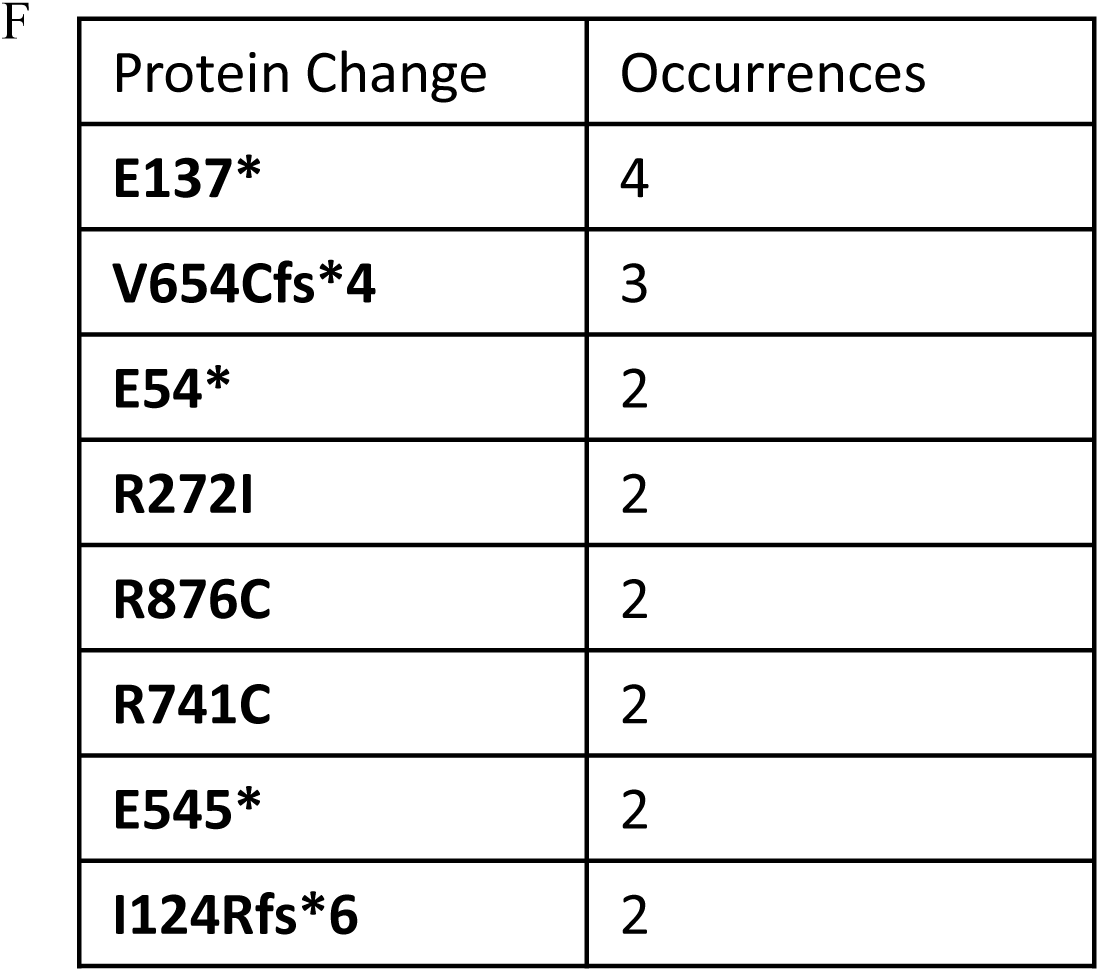
RB1 mutations in endometrial cancer. (A) Lollipop graph of RB1 mutations in endometrial cancer (cBioportal). (B) Nucleotide mutations of RB1 in endometrial cancer. (C) Types of RB1 mutations in endometrial cancer. (D) Graph for total mutation count in RB1 altered vs. unaltered endometrial cancer patients (cBioportal). (E) Graph of RB1 mutation vs. diagnosis age. (F) RB1 mutations and there occurrence in endometrial cancer.

### 3.3 Mutant RB1-E2F1 interaction

The Retinoblastoma protein (Rb) is a pivotal tumour suppressor that regulates the G1-S phase transition of the cell cycle by inhibiting E2F transcription factors, in particular E2F1 and E2F2. The conserved pocket domains of Rb1 bind E2Fs and inhibit the transcription of genes necessary for DNA synthesis. The central pocket domain of Rb1 (residues 379—792) was important for RB1-E2F1 interactions (6). As the mutations were novel, a docking study was performed to understand the interaction of Rb binding to E2F. The wild-type Rb protein established a strong and spatially coordinated contact with E2F1, mostly mediated by Rb residues Glu116, Glu117, Asp423, Glu407, His412, and Ser401(Figure S4A). These residues form direct hydrogen connections with important E2F1 regions that were rich in arginine and lysine, including Lys765, Arg908, Lys896, and Lys900. A compact and symmetrical binding was further facilitated by electrostatic bridges created by acidic residues, such as Glu280 and Glu419, as well as π–π interactions involving Tyr411 and His412 (Figure 3A), which align to the interactions shown in crystal structure 1N4M. The E137* nonsense mutation totally eliminated the structured interface. Almost all stabilising interactions seen in the wild-type complex were lost as a result of the truncation, which eliminated the whole B pocket and a sizable portion of the A pocket (Figure 3B). E2F1 could not be adequately anchored by the spatially isolated residual connections, such as those involving His406 and Glu407 (Figure S4B). Even though the R272I, it exhibited a structurally misfolded binding interface. This electrostatic environment was upset by substitution with a nonpolar isoleucine, which caused neighbouring residues to become disoriented (Figure 3C). Residues such as Glu146, His412, and Asp423 partially sustained hydrogen bonds, but their spatial orientation was severely disrupted (Figure S4C). The R741C mutant surpassed the wild-type by forming a very stable and compact complex with E2F1 (Figure 3D). Through altered polar contacts, the cysteine substitution in the B pocket appeared to form new hydrogen bonds or reinforce those already present, particularly involving His406, Glu407, Glu416, and Asp426. E2F1 residues Lys745, Lys722, and Met904 exhibit increased involvement in these interactions, forming dense bridging networks that stabilise the interface (Figure S4D). With contact sites comprising Glu416, Glu407, His412, and Asp410 loosely coordinating with E2F1 residues Lys729, Lys740, Asp718, and Lys765, interactions of R876C were more dispersed (Figure S4E). Some hydrogen bonds were still present, but unlike in the wild-type or R741C complexes, they did not form a localised interaction core (Figure 3E). The stiffness of the domain may be weakened and accurate interaction with E2F1 may be hampered if cysteine is substituted for arginine at the C-terminus, since this might disrupt local helical packing and electrostatic surface potential.

**Figure 3:**
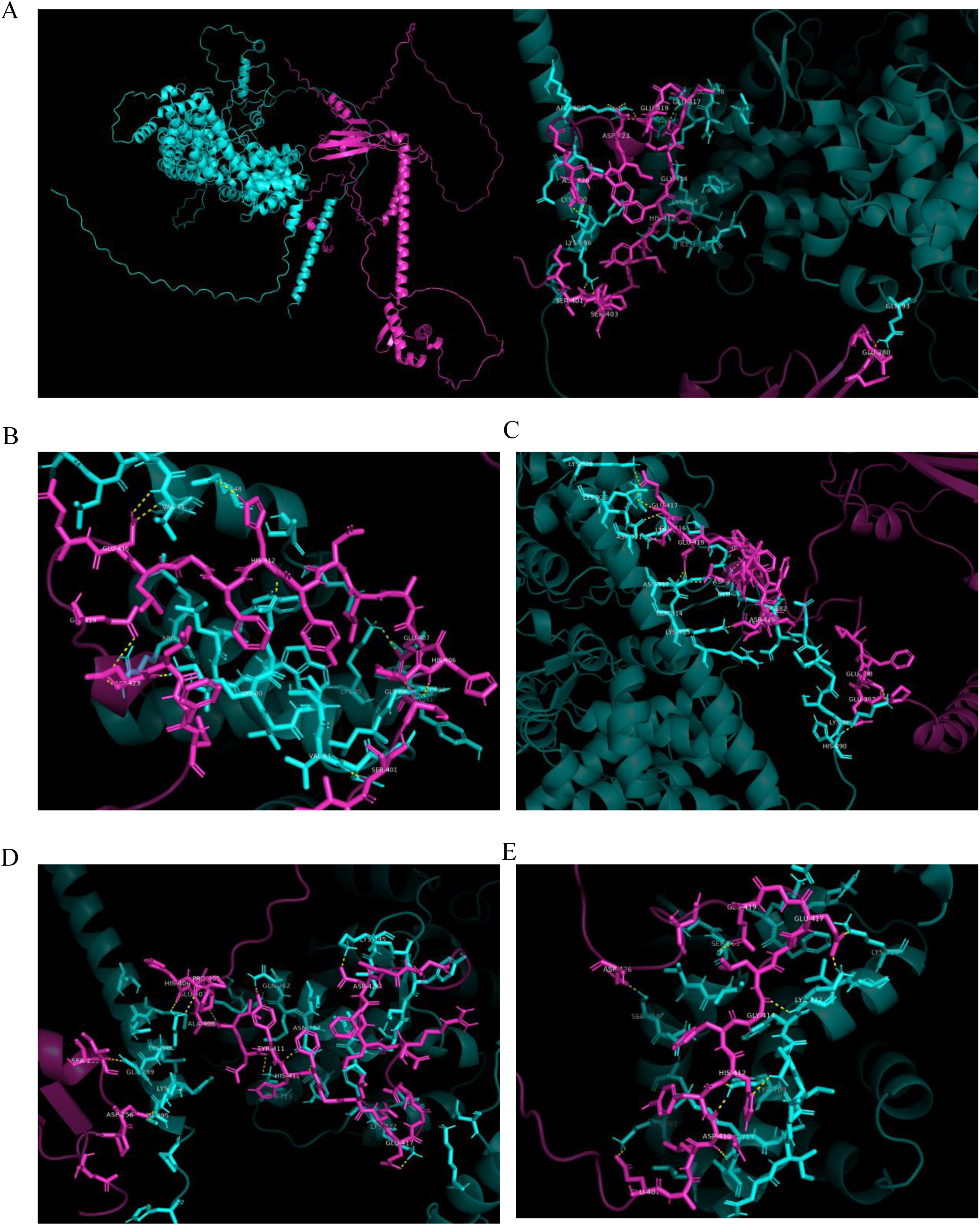
Differential binding modes of Rb mutants with E2F1. (A) Wild-type Rb (cyan) forms a compact, highly coordinated interface with E2F1 (magenta) through a network of hydrogen bonds, electrostatic bridges, and π–π interactions. (B) E137* truncation disrupts the interface by eliminating most stabilizing contacts. (C) R272I mutation causes local misfolding, disrupting key electrostatic interactions. (D) R741C mutant forms a hyper-stabilized complex via new or reinforced polar contacts. (E) R876C mutant maintains dispersed and weakened interactions, lacking a centralized binding core.

### 3.4 Mutant RB1-E2F2 interaction

Rb1 interacted with E2F2 in the wild-type complex through a broad, symmetric interface (Figure 4A). Asp433, Asp424, Asp430, Gln71, and Glu213 from Rb formed important interactions by forming a strong network of salt bridges and hydrogen bonds with Thr625, Thr408, Asn623, and Arg857 on E2F2 (Figure S5A). Just as with E2F1, the E137* binding interface was dramatically collapsed, leaving only marginal contacts intact, mostly through residues like Ser414 and Thr625 (Figure 4B). Important interface residues like Asp430, Asp433, and Gln71 were completely missing, and the resulting contact network was sparse (Figure S5B). The R272I substitution had a gain-of-binding effect due to a densely packed interaction surface with Asp343, Asp427, Asp410, Glu420, and Glu417 contributing significantly. These residues established robust hydrogen interactions with Lys94, Lys96, His129, and Lys130, a dense cluster of lysine-rich residues on E2F2 (Figure S5C). Instead of exhibiting dynamic repression, the R272I mutant seemed to establish a stiff lock between Rb and E2F2 (Figure 4C).

**Figure 4:**
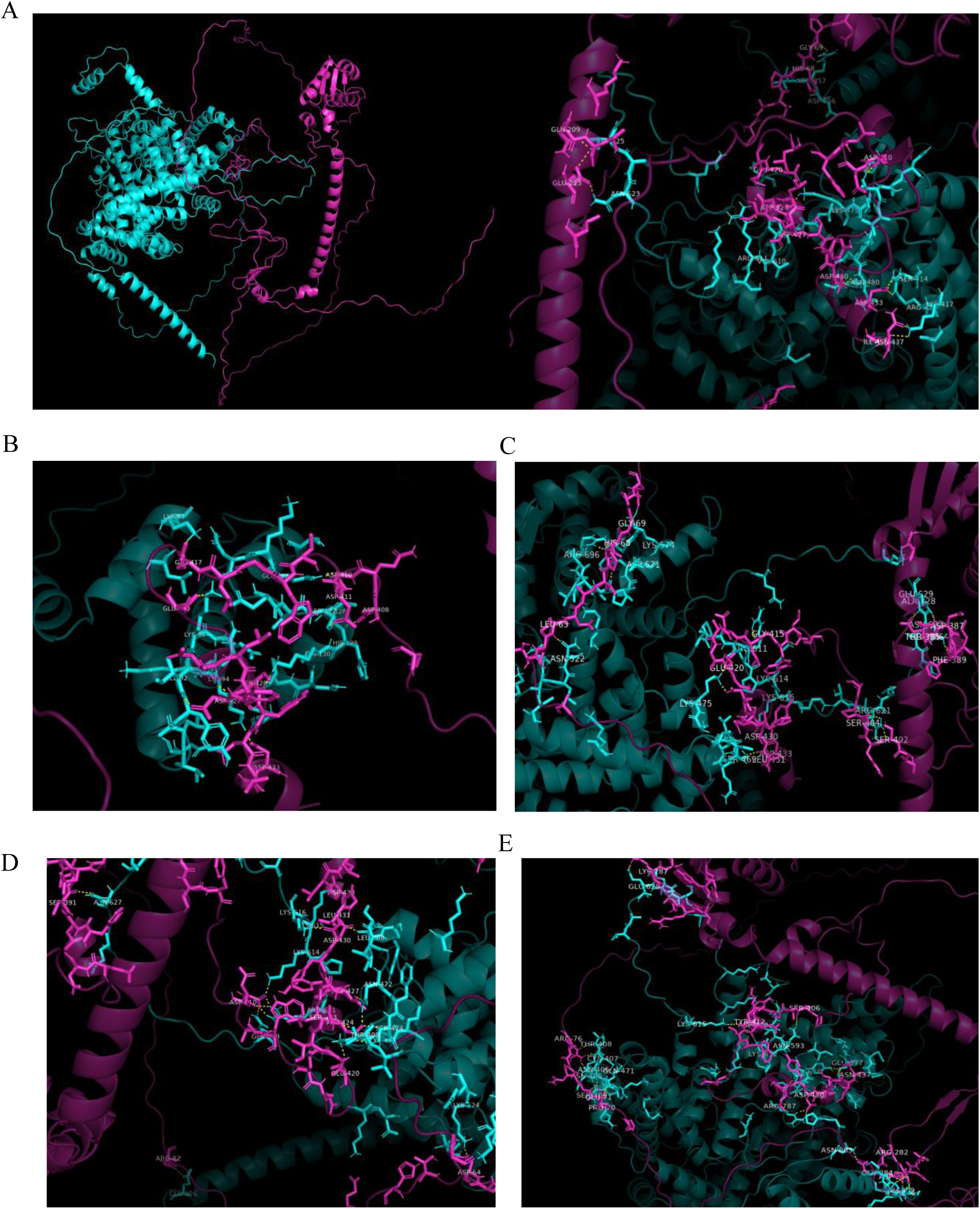
Differential binding modes of Rb mutants with E2F2. (A) Wild-type Rb (cyan) engages E2F2 (magenta) via a broad, symmetric interface dominated by acidic and polar contacts. (B) E137* mutation leads to near-complete loss of this network. (C) R272I forms a densely packed, lock-like interface through extensive hydrogen bonding with lysine-rich E2F2 regions. (D) R741C creates a rearranged but cohesive binding mode localized in the B pocket. (E) R876C establishes the most extensive and diffuse interface, involving non-canonical regions and forming a hybrid network of polar and electrostatic interactions.

The R741C mutation produced a cohesive but conformationally unique complex that was situated within the conserved B pocket of Rb. Asp430, Asp433, Glu420, Asp424, and Glu284 play a crucial role in the subtle but significant change of hydrogen bonding hotspots, which interact with Lys616, Arg611, and Asn472 of E2F2 (Figure S5D). Rearranged hydrogen bonding was notably visible in the interaction network, especially around the E2F2 recognition helix (Figure 4D). The most extensive interface seen was caused by the R876C mutation, which was located in the regulatory C-terminal region of Rb. This mutant incorporated Rb residues like Glu284, Arg282, Asp285, Ser428, and Pro70 to interact with E2F2 via a diffuse and flexible surface. On E2F2, they made contact with Arg787, Thr408, Asn405, and Lys652, forming a hybrid network of electrostatic contacts and hydrogen bonds spanning several domains (Figure S5E). The interface had a highly favourable docking score due to its tight formation and expansiveness. The structural map displayed wide contact zones over regions that were not normally engaged in canonical binding, despite the fact that the restraint violation energy was somewhat enhanced (Figure 4E).

### 3.5 Genomics analysis of RB1-altered vs. unaltered endometrial cancer patients

As the mutation count was significantly high in the RB1-altered group (Figure 2D), a genomic analysis was done, which suggested that 14,117 genes were significantly mutated in the RB1-altered population (Data sheet 4; Figure 5A). Mutations were spread across the genome, and chromosome 5 had an enriched region (Figure 5B). Pathway analysis revealed the possible cause and mechanisms of the cancer. A total of 132 significantly altered pathways were found (Data sheet 5) of which the top 50 pathways are shown (Figure 5C). The top-most mutated pathway was nicotine addiction. Out of the 40 genes of the pathway, 36 genes were mutated, which showed a strong link between nicotine addiction and RB1 mutation in endometrial cancer (Figure S6A). A large number of mutations (390/530 genes) were identified in cancer-related pathways, which include EMC receptor interaction, Wnt signalling, MAPK signalling, PI3K-Akt signalling, cAMP pathway, calcium signalling and many more (Figure S6B). More than a thousand metabolic pathway-associated genes were found to be mutated. Interestingly, the human papillomavirus infection pathway genes were also significantly mutated in the RB1 altered population (Figure S6C). Other important tumour suppressor genes (PTEN, BRCA1, BRCA2, APC, and NF1) and estrogen receptors (ESR1 and ESR2) were also co-mutated with RB1, which suggested severe consequences in the RB1-altered population (Figure 5D). As a huge number of mutations were associated with RB1 mutation, mRNA levels of were also checked in RB1 altered vs. unaltered endometrial cancer patients.

**Figure 5:**
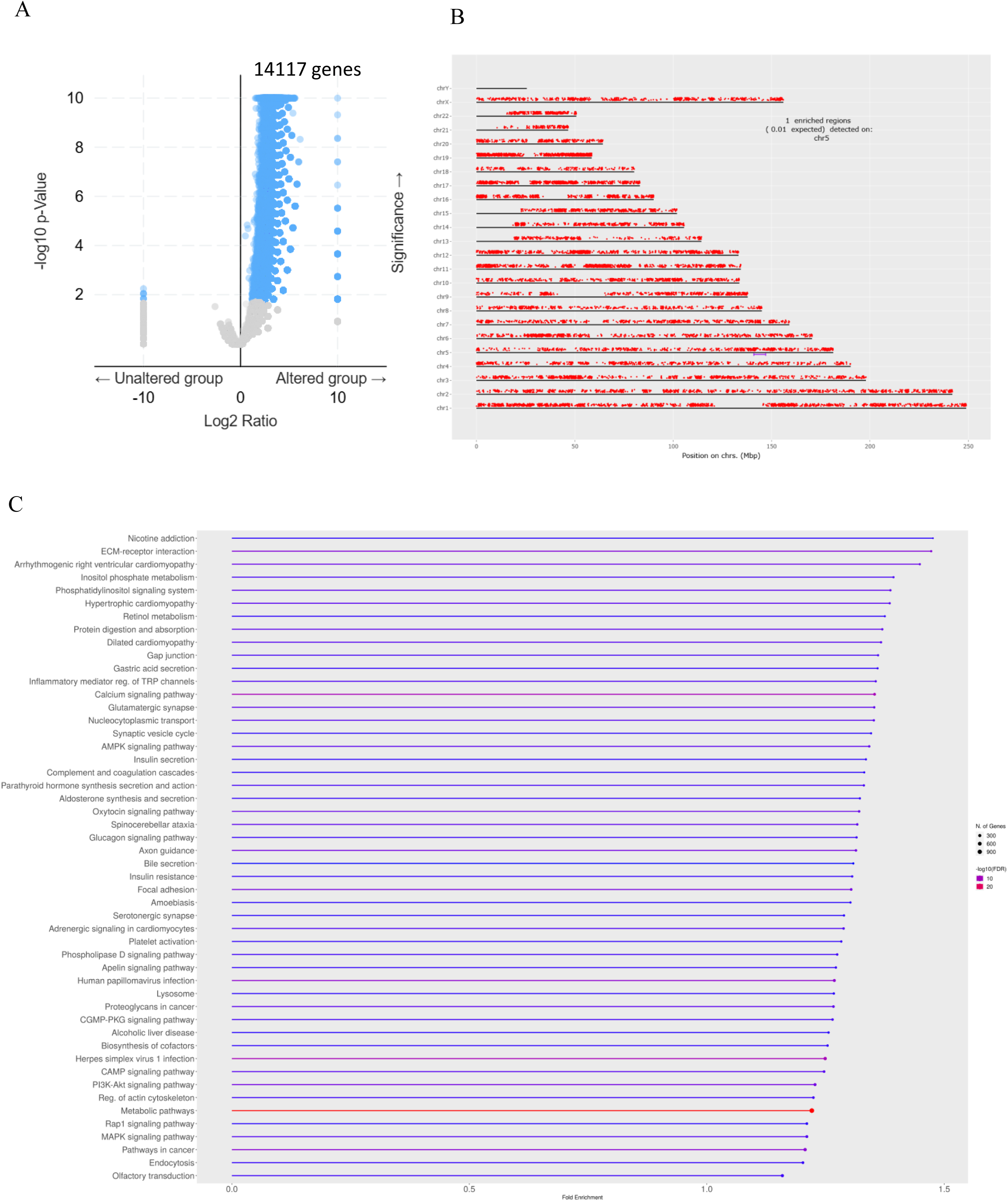
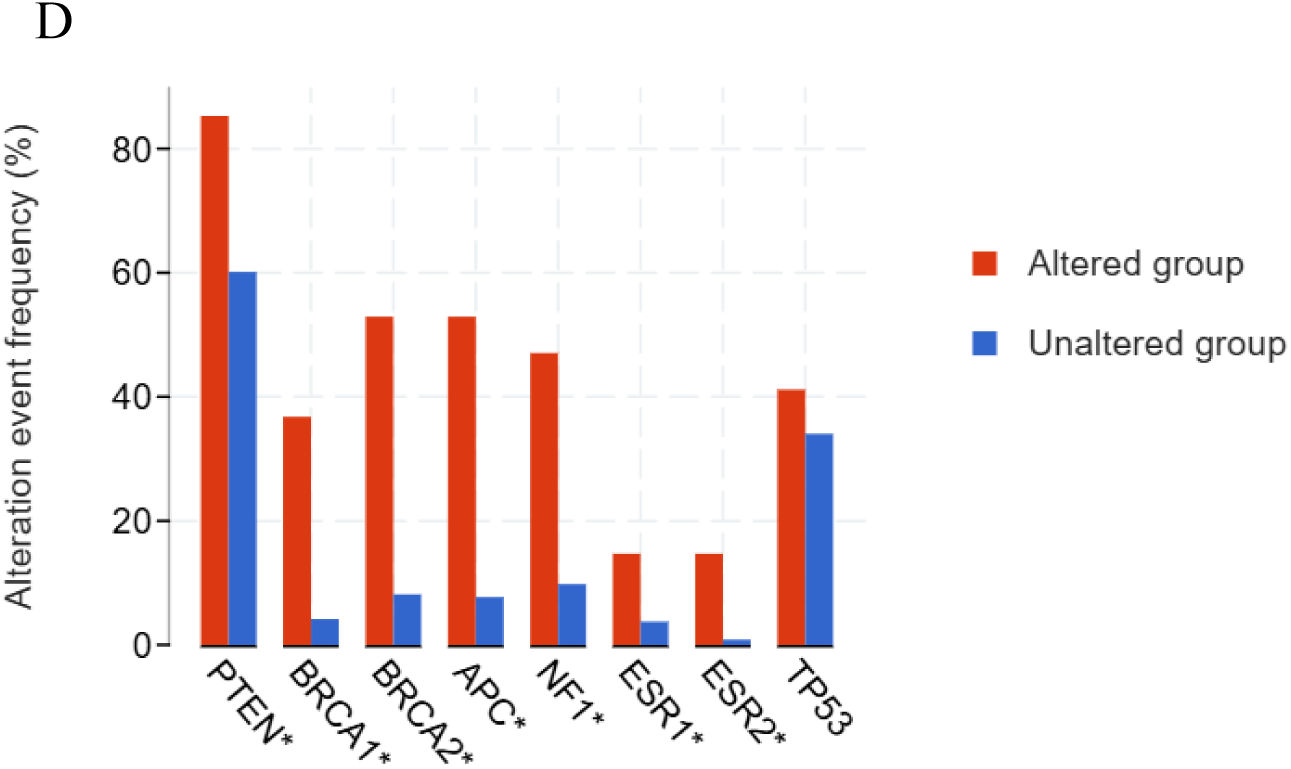
Genomics analysis of RB1 altered vs. unaltered endometrial cancer patients. (A) Volcano plot of significantly mutated genes in RB1 altered vs. unaltered endometrial cancer patients (cBioportal). (B) Positions of the mutation in the chromosomes. (C) Pathway analysis of significantly mutated genes in RB1 altered vs. unaltered endometrial cancer patients (ShinyGO). (D) The other tumour suppressor genes mutational status in RB1 altered vs. unaltered endometrial cancer patients (cBioportal).

### 3.6 Transcriptomics analysis and differentially expressed mutated gene analysis in RB1 altered vs. unaltered endometrial cancer patients

In addition to widespread mutations in the genes associated with key pathways, significant alterations in the mRNA expression levels were also observed. Transcriptomics analysis suggested that 830 genes were significantly downregulated and 4,940 genes were significantly upregulated in the RB1 altered population (Figure 6A; Data sheet 6). Most of the downregulated genes were clustered under 4 pathways (Figure 6B; Data sheet 7). The chemical carcinogenesis-reactive oxygen species and drug metabolism pathways were the most vital down-regulated pathways for cancer. The top 50 pathways of upregulated genes belonged to the DNA replication pathway, different repair pathways, the cell cycle pathway, the p53 signaling pathway, viral carcinogenesis pathways and many more (Figure 6C; Data sheet 8). The top 4 out of 5 pathways were broadly associated with DNA replication and repair, which might be the cause of the significantly higher overall mutation in the RB1 altered group (Figure 2D). As RB is a key regulator of the cell cycle, dysregulation of the cell cycle pathway genes is expected. Among the 126 genes in the pathway, 74 genes were significantly upregulated. Nuclear proliferation marker PCNA, Cyclin genes (CycD, CycE, CycA, CycH and CycB), Cyclin-dependent kinase genes (CDK1, CDK2, CDK4 and CDK6) were also ated in the RB1-altered population (Figure S7A). Genes of the p53 signaling pathway were also up-regulated. MDM2, the main negative regulator of p53, was also found to be upregulated (Figure S7B). Different viral carcinogenesis pathway genes were also upregulated, which consisted of hepatitis B virus, hepatitis C virus, Epstein-Barr virus, human papillomavirus, human T cell leukaemia virus and Kaposi’s sarcoma-associated herpesvirus-associated carcinogenesis genes (Figure S7C).

**Figure 6:**
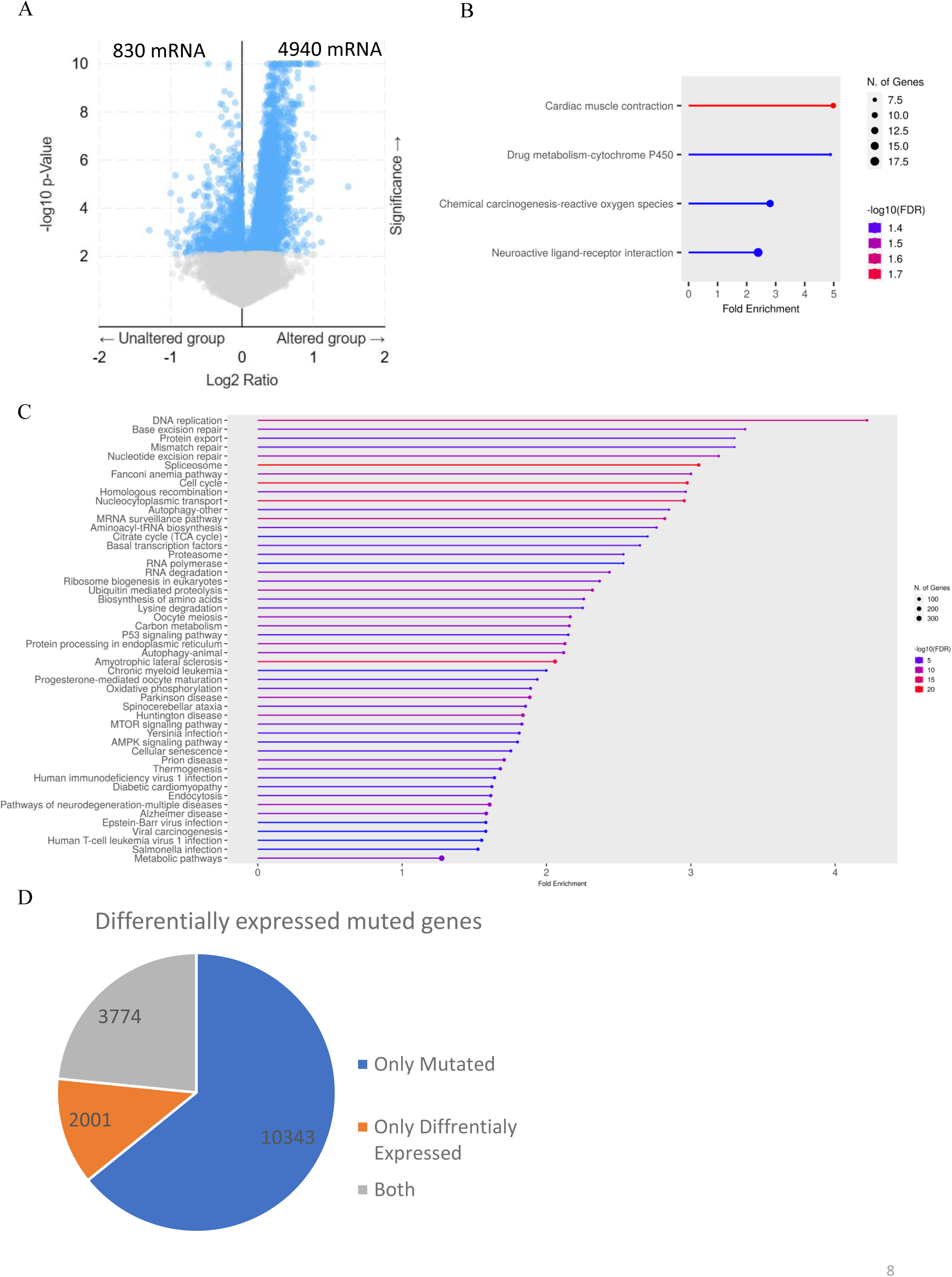
Transcriptomics analysis and differentially expressed mutated gene analysis in RB1 altered vs. unaltered endometrial cancer patients. (A) Volcano plot of differentially regulated genes in RB1 altered vs. unaltered endometrial cancer patients (cBioportal). (B) Pathway analysis of significantly downregulated genes in RB1 altered vs. unaltered endometrial cancer patients (ShinyGO). (C) Pathway analysis of significantly upregulated genes in RB1 altered vs. unaltered endometrial cancer patients (ShinyGO). (D) Pie chart of differentially expressed mutated genes. (C) Bar graph of upregulated genes mutated involved in different pathways.

A total of 65.35% of differentially expressed genes were also mutated (Figure 6D, Data sheet 9). Most of the differentially regulated genes present in different pathways were mutated in the RB1-altered population. 77.7% of DNA replication pathways were differentially regulated and mutated. Five out of the top six pathways of differentially expressed and mutated genes were related to replication, repair and recombination pathways, which might be the reason behind 14,117 co-mutated genes in RB1 populations. The cancer-related pathways (mTOR signalling, p53 signalling, etc.) genes were also represented. Additionally, 45% of the genes in the cell cycle pathways were both differentially regulated and mutated. Human papillomavirus infection-associated and metabolic pathway genes were also dysregulated at both genomic and transcriptomic levels (Figure S8; Data sheet 10). These findings are particularly unique to endometrial cancer and were checked in another endometrial cancer dataset.

### 3.7 Checking of key observations in an endometrial cancer dataset and other gynaecological cancers

1,315 endometrial cancer samples were taken from the MSK MetTropism (MSK, Cell 2021) dataset, which did not have any overlap with the samples of the previously used endometrial cancer dataset (Endometrial Carcinoma (TCGA, GDC). In this cohort, 383 genes were significantly found to be co-mutated with RB1 (Figure 7A; Data sheet 11). Pathway analysis revealed that these mutated genes were involved in repair pathways (mismatch repair, nucleotide excision repair and base excision repair), cancer-related pathways (p53 signalling pathway, mTOR signalling pathway, HIF1 signalling pathway, etc.), cell cycle pathways, metabolic pathways and Human papillomavirus infection pathways, which is consistent with the previous observations (5C and 6C). Other essential tumour suppressor genes (PTEN, BRCA1, BRCA2, APC, and NF1) were also significantly mutated in the RB1-altered patients. The estrogen receptor gene, ESR1, was also found to be co-mutated with RB1 (Figure 7B). The G→T nucleotide mutation and missense mutations were the most commonly found mutations in RB1 (Figure 7C and 7D). Rb1-altered patients exhibited a significantly higher overall mutation burden (Figure 7E). Moreover, a higher frequency of Rb1 mutations was observed among the endometrial cancer patients of the age group between 40-50 years (Figure 7F; Data sheet 14).

**Figure 7:**
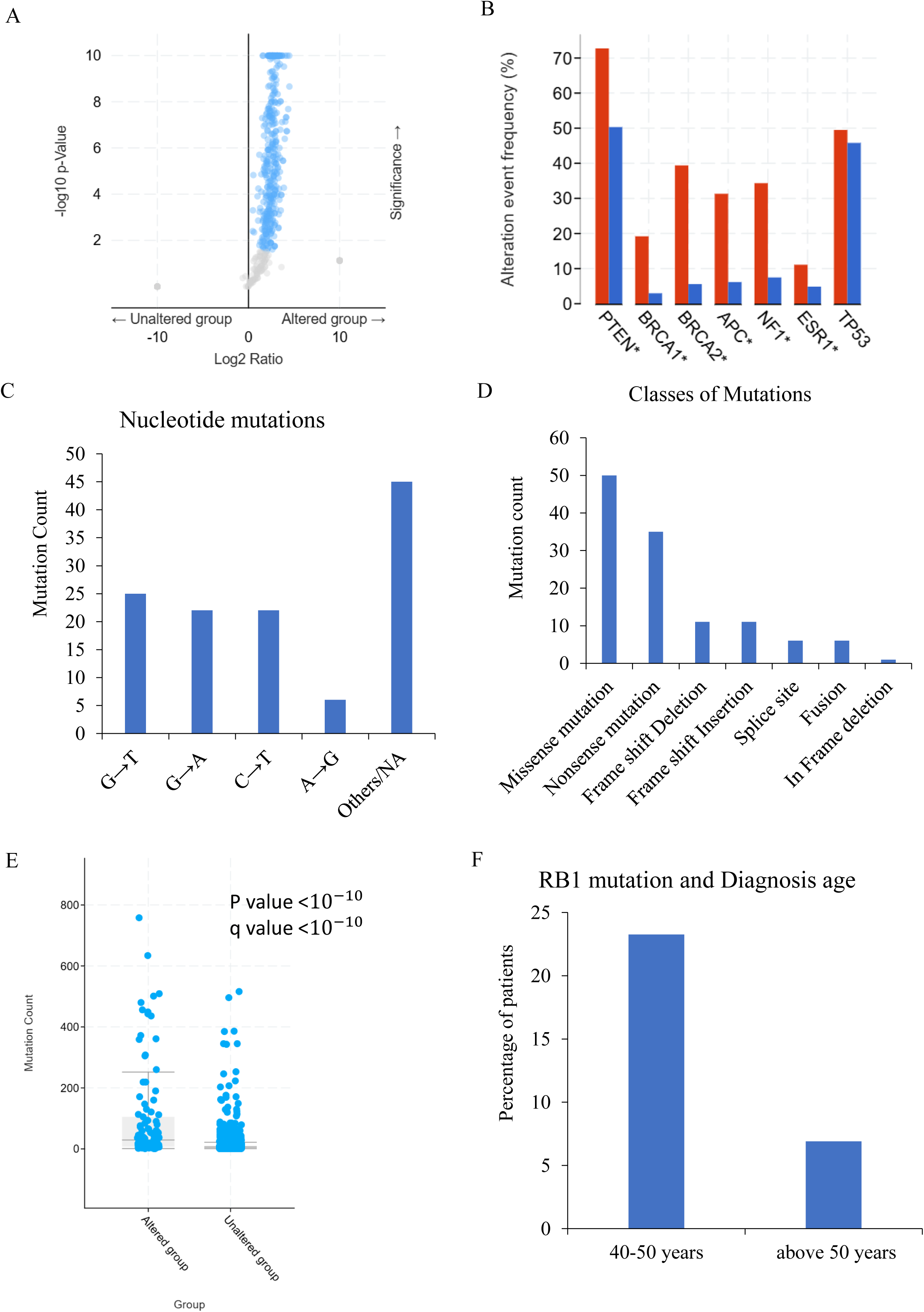
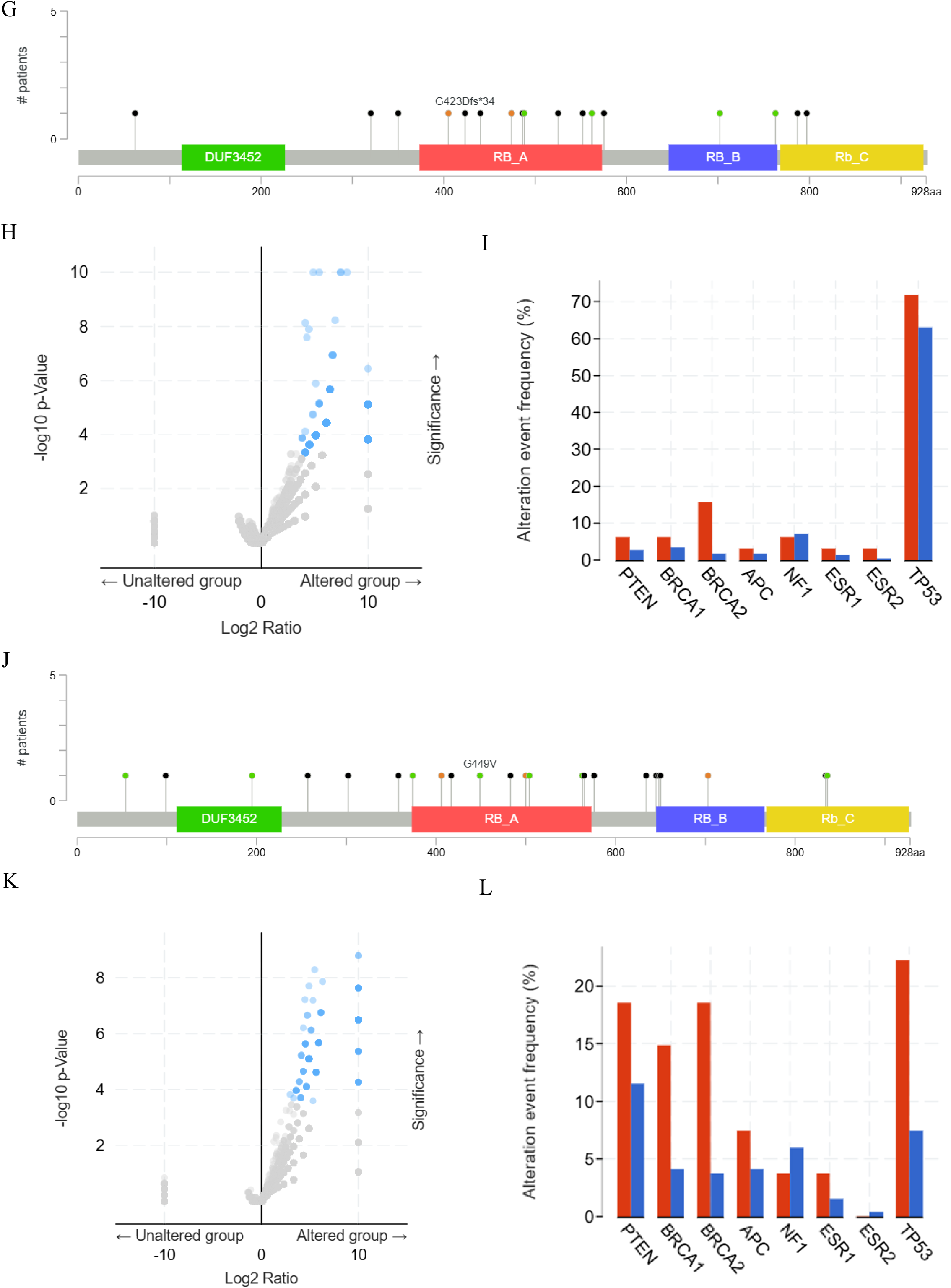
Checking of key observations in an independent endometrial cancer dataset and other gynaecological cancers. (A) Volcano plot of differentially regulated genes in RB1 altered vs. unaltered endometrial cancer patients (cBioportal). (B) Other tumour suppressor gene’s status in RB1 altered vs. unaltered in another endometrial cancer patients (cBioportal). (C) Nucleotide mutations of RB1 in another endometrial cancer dataset. (D) Types of RB1 mutations in another endometrial cancer dataset. (E) Graph for total mutation count in RB1 altered vs. unaltered another endometrial cancer patients (cBioportal). (F) Graph of RB1 mutation vs. diagnosis age validation. (G) Lollipop graph of RB1 mutations in ovarian cancer (cBioportal). (H) Volcano plot of differentially regulated genes in RB1 altered vs. unaltered ovarian cancer patients (cBioportal). (I) The other tumour suppressor gene mutational status in RB1-altered versus unaltered ovarian cancer patients (cBioportal). (J) Lollipop graph of RB1 mutations in cervical cancer (cBioportal). (K) Volcano plot of differentially regulated genes in RB1 altered vs. unaltered cervical cancer patients (cBioportal). (L) The other tumour suppressor gene mutational status in RB1-altered versus unaltered cervical cancer patients (cBioportal).

The RB1 mutations were also checked in other gynaecological cancers (ovarian cancer and cervical cancer). Only 6% RB1 mutation was found in 602 samples of the ovarian cancer dataset (Figure 7G). 322 genes were co-mutated with RB1, but there was no significant enrichment found in the pathway analysis (Figure 7H and Data sheet 12). Cervical cancer datasets (297 samples) had 9% RB1 mutations, and only 139 genes were co-mutated with RB1(Figure 7J and 7K; Data sheet 13). There was no significant enrichment in the pathway analysis of these 139 co-mutated genes. Other essential tumour suppressor genes (PTEN, BRCA1, BRCA2, APC, and NF1) and estrogen receptors (ESR1 and ESR2) were not significantly mutated in the RB1-altered patients (Figure 7I and 7L). These results suggest that the findings in endometrial cancer patients were very unique when compared to other gynaecological cancers.

## 4. Discussion

RB1 ranks fourth among the tumour suppressor genes in terms of frequency of homozygous loss (8). RB1 mutations in retinoblastoma, bladder cancer, lung cancer, breast cancer, and prostate cancer are well known (13–16,23). The RB1 mutations and significant upregulation of RB1 protein were found in endometrial cancer (Figure 1C,1D, and S3C). Many genes (PTEN, PIK3CA, PIK3R1, CTNNB1, ARID1A, K-RAS, CTCF, RPL22, TP53, FGFR2, and ARID5B) were known to be mutated in endometrial cancer (17). A systematic study of the RB1 mutation in endometrial cancer was done for the first time in the current study. The mutation frequency of RB1 was 12%, which was lower than the RB1 mutation in retinoblastoma (Figure 2A)(24). Previous reports on other cancers have studied the correlation between age and RB1 mutation, but the relationship remained unclear (25,26). RB1 mutation frequencies were high in endometrial cancer patients under 50 years (Figure 2E). This indicates that endometrial cancer patients before menopause have higher chances of RB1 mutations and associated co-mutations in other tumour suppressor genes and estrogen receptor genes, which could increase the severity of endometrial cancer. (figure 5D). ESR1 and ESR2 polymorphisms and mutations were associated with causing regulatory changes in endometrial cancer(27,28). These findings were also confirmed using a different endometrial cancer dataset (Figure 7B-F).

Previous literature has established RB1 as a tumor suppressor that inhibits E2F1/E2F2 through structured pocket domain interactions (29,30), but the effects of many RB1 mutations remain uncharacterized. In this study, RB1-E2F1 and RB1-E2F2 interactions were analyzed through molecular docking to assess the impact of novel RB1 mutations. Wild-type RB1 formed stable, structured complexes with both E2F1 and E2F2 via key acidic residues. Mutations like E137* caused severe loss of interaction due to truncation, while R272I and R741C altered or enhanced binding through misfolded or reinforced interfaces. R876C showed dispersed interactions with potential structural instability. These findings reveal mutation-specific effects on E2F binding. However, functional assays and in vivo validation are still needed to confirm how these altered interactions influence cell cycle regulation and tumorigenesis.

Retinoblastoma genes also regulate genome stability (31). The 14,117 genes were significantly mutated with RB1 mutations in endometrial cancer and the replication and repair machinery genes were differentially regulated and highly mutated (Figure 5A, 5C, 6C and S8). Genes involved in the Human papillomavirus infection pathway were found to be both mutated and differentially regulated in RB1-altered endometrial cancer samples (Figure 5C, 6C and S8).

Although Human papillomavirus has a limited role in causing endometrial cancer (32), the RB1 mutations in Human papillomavirus-associated cervical cancer are well studied (33). This raises the possibility that the HPV vaccination could potentially offer preventive benefits not only for cervical cancer but also for a subset of endometrial cancers. These findings suggest that Human papillomavirus infection can be a cause of RB1 mutation-associated endometrial cancer. RB1 protein can bind with E2F and inhibit the transactivation of cell cycle-regulating genes (7,34). In endometrial cancer, RB1-mutated populations are also shown to have differentially regulated and mutated Cyclin genes and Cyclin-dependent kinase genes (Figure 5C, 6C and S8). The RB-E2F transcription machinery is known to regulate the different metabolic pathways like TCA cycle, OXPHOS, glucose oxidation (PDK4), fatty acid oxidation (FAO), nucleotide biosynthesis, and glutamine metabolism (ASCT2, GLS1)(35). Metabolic reprogramming and cancer-related pathway dysfunction were also found in the RB1-altered population (Figure 5C, 6C and S8). Most of the pathways were also analysed in the endometrial cancer dataset of MetTropism, and the results remain the same. These findings were unique and confined only to endometrial cancer and not present in other gynaecological cancers (7G-L). This resulted in the mutation and differential expression of genes involved in the cell cycle, metabolic pathways and cancer related pathways, ultimately contributing to the regulation and progression of endometrial cancer (Figure 8).

**Figure 8:**
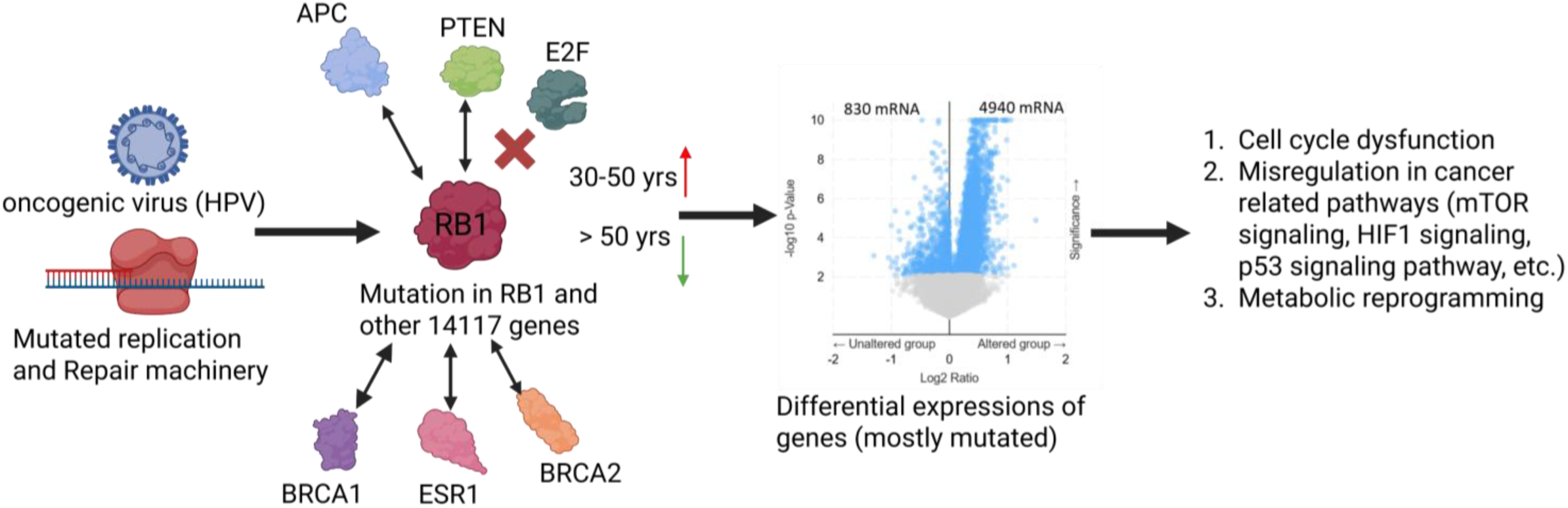
RB1 gene mutations and their association with Endometrial Cancer. Due to mutations in the replication-repair machinery and viral infection (HPV), 14117 other genes were mutated, along with RB1. RB1 and E2F interaction was impaired. 5770 genes involved in cell cycle regulation, cancer-related pathways, and metabolic reprogramming were expressed differentially and regulated the endometrial cancer.

Our findings highlight RB1 not only as a tumor suppressor via E2F regulation but also as a central mediator in the metabolic, viral response, and cell cycle networks of endometrial tumors. These findings may have diagnostic and therapeutic implications, particularly in guiding targeted therapies for RB1-mutated patients or stratifying younger patients for closer surveillance. These insights pave the way for further investigations into RB1-directed therapies and open new questions regarding viral and metabolic interplay in endometrial tumorigenesis.

## Supporting information

https://drive.google.com/drive/folders/1hTJMum6_PKdYev6cjZhwpA0QL_rXhmk7?usp=sharing.

## Acknowledgements

DBT is acknowledged for the JRF fellowship to PKG. Dr. Karyala Prashanthi was acknowledged for her valuable suggestions.

## 5. Author contribution

Pritam kr. Ghosh and Saumitra Das: Conception and design of studies-analysis, and interpretation. Pritam kr. Ghosh, Somrita Das, Gagan Gaurav, Sahana Ghosh, Arindam Maitra, Manjula Das and Saumitra Das: Interpretation of results, article writing, and article editing. Saumitra Das: Funding acquisition, supervision and project management.

## 7. Data availability statement

No original data was generated. All mutations and mRNA level data were taken from cBioportal (https://www.cbioportal.org/datasets). Data sheets of the analysed data are available at https://drive.google.com/drive/folders/1hTJMum6_PKdYev6cjZhwpA0QL_rXhmk7?usp=sharing.

## 8. Abbreviations

RB1: Retinoblastoma gene
pRB: retinoblastoma protein
E2F: Early region 2 binding factor
HPV: Human Papillomavirus
FDR: false discovery rate
UCES: Uterine corpus endometrial carcinoma

## Supplementary figure Legends

**Figure S1: Patient sample information**

(A) Pei chart of cancer types (MSK MetTropism dataset). (B) Pei chart of the Race category (MSK MetTropism dataset). (C) Pei chart of Race category (Endometrial Carcinoma (TCGA, GDC) https://www.cbioportal.org/study/summary?id=ucec_tcga_gdc). (D) Pei chart of the Race category Ovarian Cancer (TCGA, GDC; https://www.cbioportal.org/study/summary?id=hgsoc_tcga_gdc). (E) Pei chart of the Race category Cervical Squamous Cell Carcinoma (TCGA, PanCancer Atlas; https://www.cbioportal.org/study/summary?id=cesc_tcga_pan_can_atlas_2018),

**Figure S2: RB1 mutation analysis in Retinoblastoma**

(A) Types of RB1 mutations in retinoblastoma. (B) Nucleotide mutations of RB1 in retinoblastoma.

**Figure S3: RB1 Expressions in normal tissues**

(A) RB1 mRNA expression across tissues (human protein atlas). (B) RB1 protein expression across tissues (human protein atlas). (C) Protein expression of RB1 in uterine corpus endometrial carcinoma (UALCAN).

**Figure S4. Comparative interaction profiles of wild-type and mutant Rb proteins with E2F1.**

(A) Wild-type Rb (WT Rb) interacting with E2F1: The docking interaction map reveals a dense network of hydrogen bonds (green dashed lines) between conserved residues of Rb and E2F1, notably involving Glu416, Glu419, Asp423, and His412 from Rb interacting with Lys860, Arg866, and Lys900 from E2F1. Hydrophobic interactions (red arcs) are prominently observed around His412 and Leu409, supporting a stable binding interface.

(B) E137Stop Rb mutant with E2F1: This premature stop codon truncation leads to a substantial reduction in the number of hydrogen bonds and a shift in the binding interface. The loss of the C-terminal domain results in fewer stabilizing contacts, with only partial compensation by residues such as Glu79, Arg723, and Tyr101, indicating a compromised binding affinity.

(C) R272I Rb mutant with E2F1: Substitution of a positively charged arginine with a non-polar isoleucine results in loss of key electrostatic interactions. Although some hydrogen bonding is retained via Asp423 and Glu419, there is a noticeable shift toward hydrophobic stabilization. The overall interaction profile suggests moderate affinity, with altered contact residues including Ala917 and Glu880.

(D) R742C Rb mutant with E2F1: Mutation of Arg742 to cysteine slightly modifies the charge distribution while preserving several key polar interactions. Hydrogen bonds between Glu410, Glu416, and Asp426 of Rb and Lys740 and Ser758 of E2F1 are maintained. Hydrophobic interactions are concentrated around Lys745 and Met804, suggesting a moderate degree of interface preservation.

(E) R876C Rb mutant with E2F1: Despite the arginine-to-cysteine substitution, this mutant retains the most interaction features similar to WT. Extensive hydrogen bonding is observed involving Glu410, Glu417, and His412, with multiple E2F1 lysines (Lys765, Lys780, Lys713) engaging in polar and hydrophobic interactions. This suggests a relatively stable binding interface among the tested mutants.

Colour coding: Hydrogen bonds are indicated by green dashed lines, hydrophobic contacts by red semicircles, and interacting residues are labeled with amino acid and chain identifiers. E2F1 residues are typically marked as chain A, and Rb residues as chain B.

**Figure S5. Comparative interaction profiles of wild-type and mutant Rb proteins with E2F2.**

(A) Wild-type Rb (WT Rb) interacting with E2F2: The interaction map displays a dual anchoring architecture with dense hydrogen bonding between acidic Rb residues (e.g., Asp343, Asp410, Asp424, Glu209) and basic E2F2 residues (e.g., Arg418, Arg611, Lys475). Hydrophobic contacts involving Leu468 and Tyr615 support a stable, cooperative binding interface.

(B) E137Stop Rb mutant with E2F2: Truncation of the B pocket leads to significant loss of hydrogen bonds and interaction density. Only fragmented contacts involving Glu407, Asp427, and His129 remain, lacking a central interaction core and resulting in a collapsed interface.

(C) R272I Rb mutant with E2F2: The Arg to Ile substitution disrupts the electrostatic network, shifting the interface geometry. Although bonds involving Asp343, Glu420, and Asp430 are retained, contacts appear scattered and less cooperative. Isolated π-interaction patches indicate structural misalignment despite moderate electrostatic energy.

(D) R741C Rb mutant with E2F2: This mutant frms a highly compact and stabilized interface. Hydrogen bonds involving Asp410, Asp424, and Asp433 anchor key E2F2 residues (Lys574, Tyr615, Arg611). Enhanced hydrophobic clustering around Leu468 and π-interactions contribute to a gain-of-binding phenotype.

(E) R876C Rb mutant with E2F2: The interface is broad but disorganized. Hydrogen bonds are present but diffuse, involving Ser428, Arg282, and Asp430 with the C-terminal region of E2F2. Structural dispersion and C-terminal misalignment likely reduce binding efficiency.

Colour coding: Hydrogen bonds are indicated by green dashed lines, hydrophobic contacts by red semicircles, and interacting residues are labeled with amino acid and chain identifiers. E2F2 residues are typically chain A, and Rb residues chain B.

**Figure S6: Pathways with a high percentage of mutated genes.**

(A) Nicotine addiction pathway. (B) Pathways in cancer. (C) Human papillomavirus infections. Red coloured genes in the pathway are mutated genes.

**Figure S7: Upregulated genes pathway**

(A) Cell cycle pathway. (B) p53 gene pathway. (C) Viral carcinogenesis. Upregulated genes are marked in red colour.

**Figure S8: Bar graph of differentially regulated mutated genes involved in different pathways.**

**Table 1: (A) Interactions of Rb and its mutants with E2F1. (B) Interactions of Rb and its mutants with E2F2.**

