## Supplementary material for "RB1 gene mutations and their association with Endometrial Cancer": https://drive.google.com/drive/folders/1hTJMum6_PKdYev6cjZhwpA0QL_rXhmk7?usp=sharing.

S1.

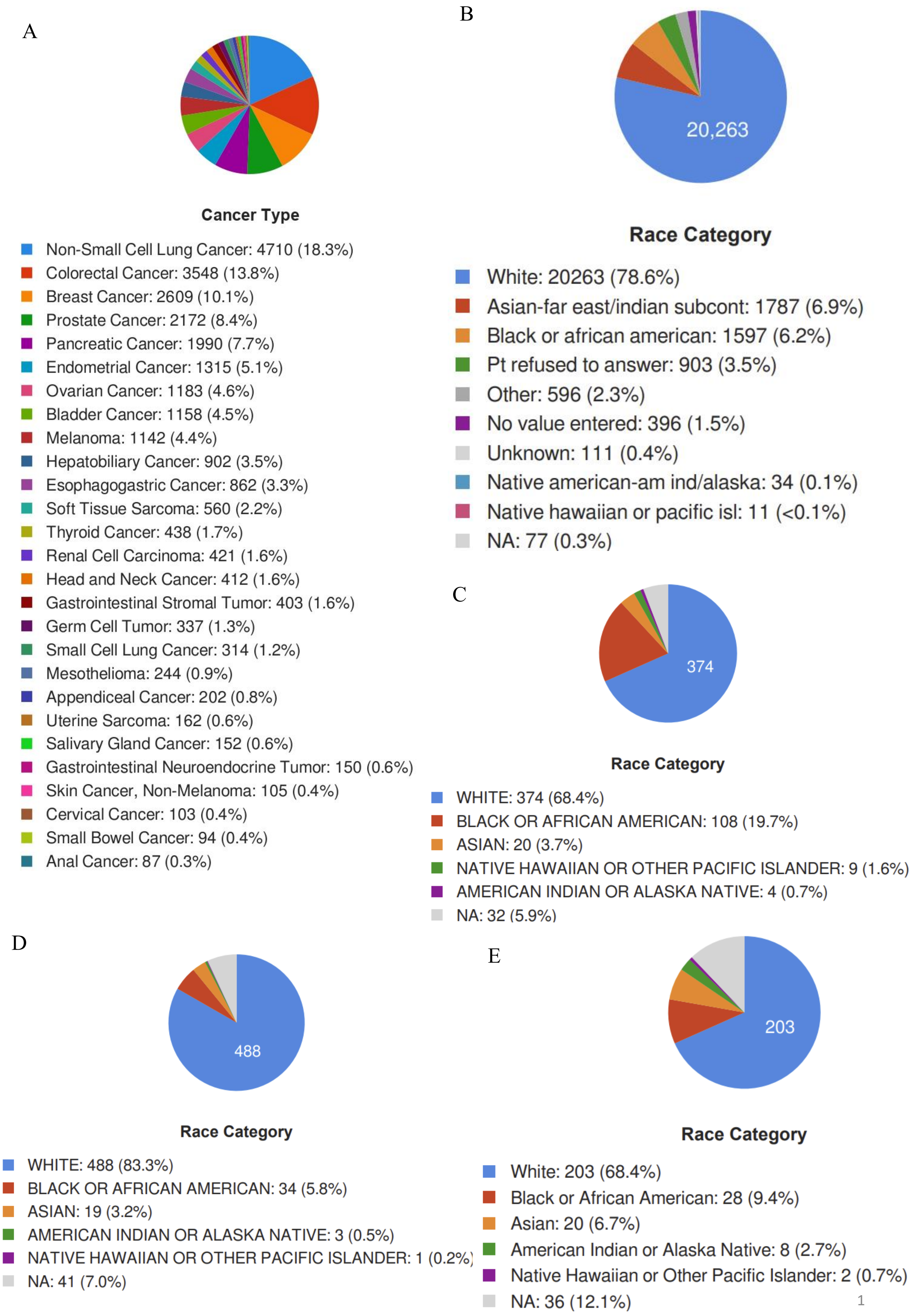
A
Cancer Type
Non-Small Cell Lung Cancer: 4710 (18.3%)
Colorectal Cancer: 3548 (13.8%)
Breast Cancer: 2609 (10.1%)
Prostate Cancer: 2172 (8.4%)
Pancreatic Cancer: 1990 (7.7%)
Endometrial Cancer: 1315 (5.1%)
Ovarian Cancer: 1183 (4.6%)
Bladder Cancer: 1158 (4.5%)
Melanoma: 1142 (4.4%)
Hepatobiliary Cancer: 902 (3.5%)
Esophagogastric Cancer: 862 (3.3%)
Soft Tissue Sarcoma: 560 (2.2%)
Thyroid Cancer: 438 (1.7%)
Renal Cell Carcinoma: 421 (1.6%)
Head and Neck Cancer: 412 (1.6%)
Gastrointestinal Stromal Tumor: 403 (1.6%)
Germ Cell Tumor: 337 (1.3%)
Small Cell Lung Cancer: 314 (1.2%)
Mesothelioma: 244 (0.9%)
Appendiceal Cancer: 202 (0.8%)
Uterine Sarcoma: 162 (0.6%)
Salivary Gland Cancer: 152 (0.6%)
Gastrointestinal Neuroendocrine Tumor: 150 (0.6%)
Skin Cancer, Non-Melanoma: 105 (0.4%)
Cervical Cancer: 103 (0.4%)
Small Bowel Cancer: 94 (0.4%)
Anal Cancer: 87 (0.3%)
B
20,263
Race Category
White: 20263 (78.6%)
Asian-far east/indian subcont: 1787 (6.9%)
Black or african american: 1597 (6.2%)
Pt refused to answer: 903 (3.5%)
Other: 596 (2.3%)
No value entered: 396 (1.5%)
Unknown: 111 (0.4%)
Native american-am ind/alaska: 34 (0.1%)
Native hawaiian or pacific isl: 11 (<0.1%)
NA: 77 (0.3%)
C
374
Race Category
WHITE: 374 (68.4%)
BLACK OR AFRICAN AMERICAN: 108 (19.7%)
ASIAN: 20 (3.7%)
NATIVE HAWAIIAN OR OTHER PACIFIC ISLANDER: 9 (1.6%)
AMERICAN INDIAN OR ALASKA NATIVE: 4 (0.7%)
NA: 32 (5.9%)
D
488
Race Category
WHITE: 488 (83.3%)
BLACK OR AFRICAN AMERICAN: 34 (5.8%)
ASIAN: 19 (3.2%)
AMERICAN INDIAN OR ALASKA NATIVE: 3 (0.5%)
NATIVE HAWAIIAN OR OTHER PACIFIC ISLANDER: 1 (0.2%)
NA: 41 (7.0%)
E
203
Race Category
White: 203 (68.4%)
Black or African American: 28 (9.4%)
Asian: 20 (6.7%)
American Indian or Alaska Native: 8 (2.7%)
Native Hawaiian or Other Pacific Islander: 2 (0.7%)
NA: 36 (12.1%)

S2.

A

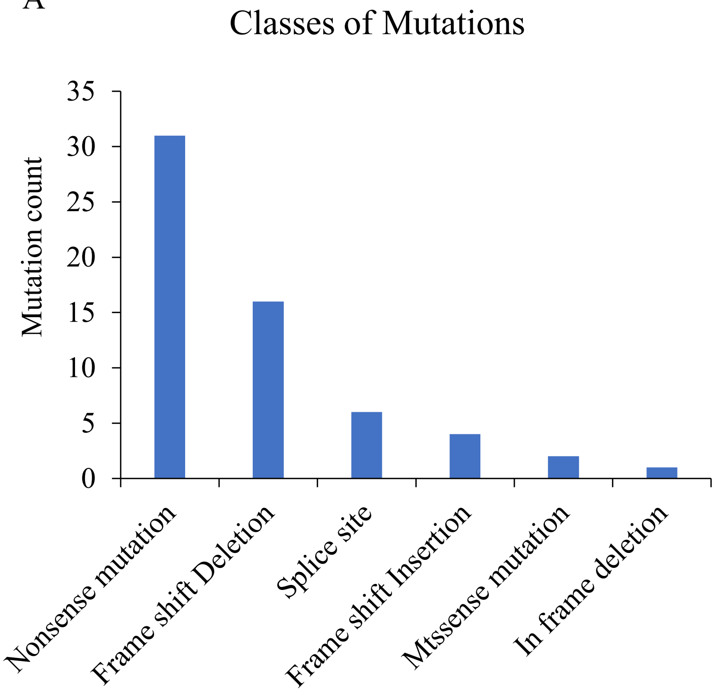
Classes of Mutations
Mutation count
35
30
25
20
15
10
5
0
Nonsense mutation
Frame shift Deletion
Splice site
Frame shift Insertion
Mtssense mutation
In frame deletion

B

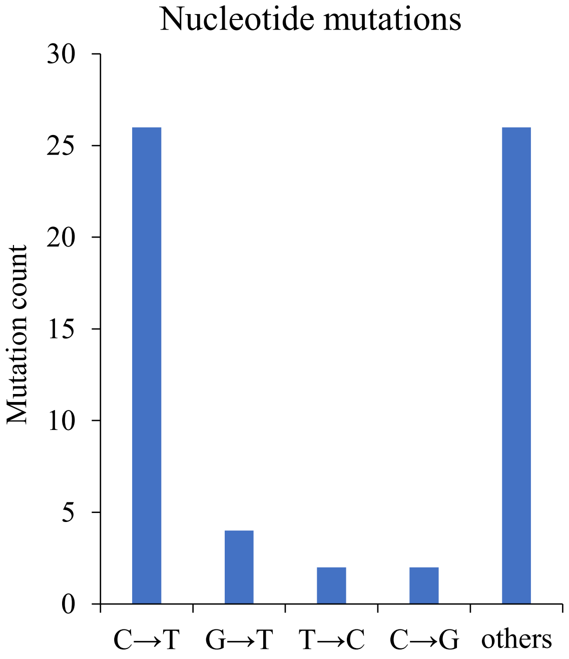
Nucleotide mutations
Mutation count
30
25
20
15
10
5
0
C→T
G→T
T→C
C→G
others

S3.

A

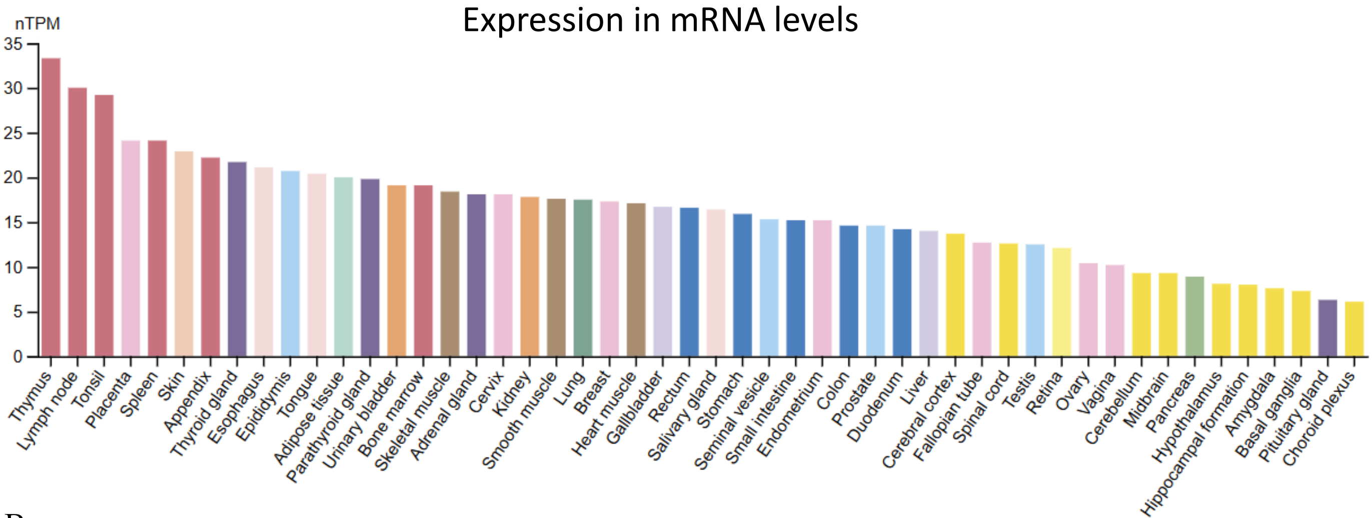
Expression in mRNA levels
nTPM
35
30
25
20
15
10
5
0
Thymus
Lymph node
Tonsil
Placenta
Spleen
Skin
Appendix
Thyroid gland
Esophagus
Epididymis
Tongue
Adipose tissue
Parathyroid gland
Urinary bladder
Bone marrow
Skeletal muscle
Adrenal gland
Cervix
Kidney
Smooth muscle
Lung
Breast
Heart muscle
Gallbladder
Rectum
Salivary gland
Stomach
Seminal vesicle
Small intestine
Endometrium
Colon
Prostate
Duodenum
Liver
Cerebral cortex
Fallopian tube
Spinal cord
Testis
Retina
Ovary
Vagina
Cerebellum
Midbrain
Pancreas
Hypothalamus
Hippocampal formation
Amygdala
Basal ganglia
Pituitary gland
Choroid plexus

B

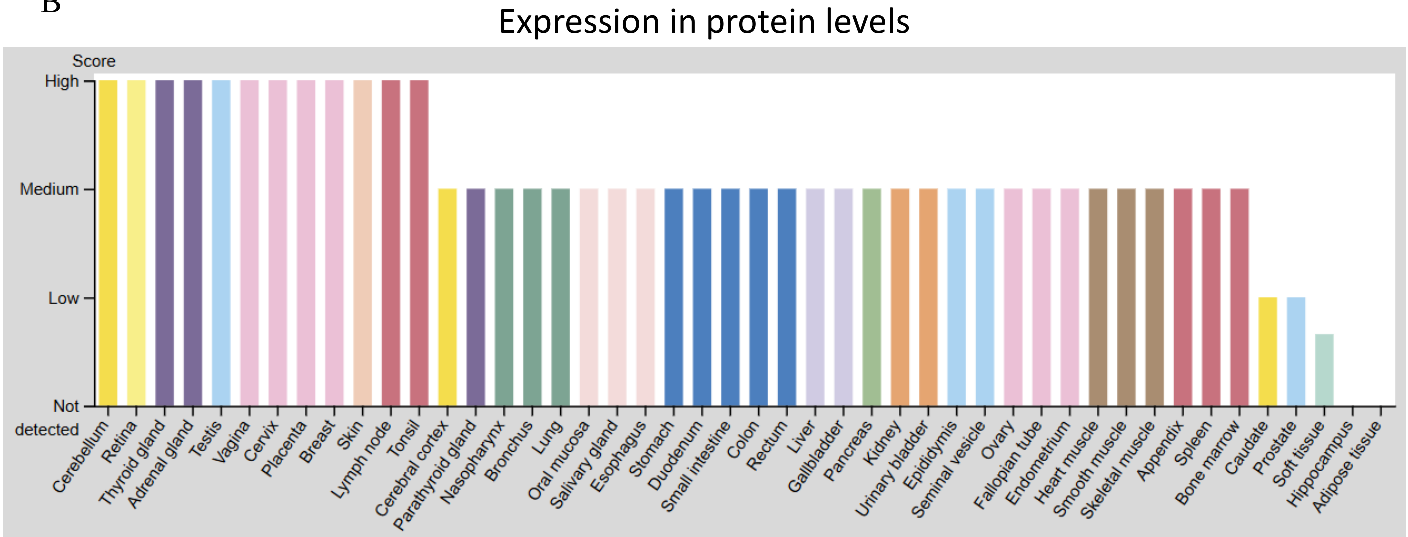
Expression in protein levels
Score
High
Medium
Low
Not detected
Cerebellum
Retina
Thyroid gland
Adrenal gland
Testis
Vagina
Cervix
Placenta
Breast
Skin
Lymph node
Tonsil
Cerebral cortex
Parathyroid gland
Nasopharynx
Bronchus
Lung
Oral mucosa
Salivary gland
Esophagus
Stomach
Duodenum
Small intestine
Colon
Rectum
Liver
Gallbladder
Pancreas
Kidney
Urinary bladder
Epididymis
Seminal vesicle
Ovary
Fallopian tube
Endometrium
Heart muscle
Smooth muscle
Skeletal muscle
Appendix
Spleen
Bone marrow
Caudate
Prostate
Soft tissue
Hippocampus
Adipose tissue

C

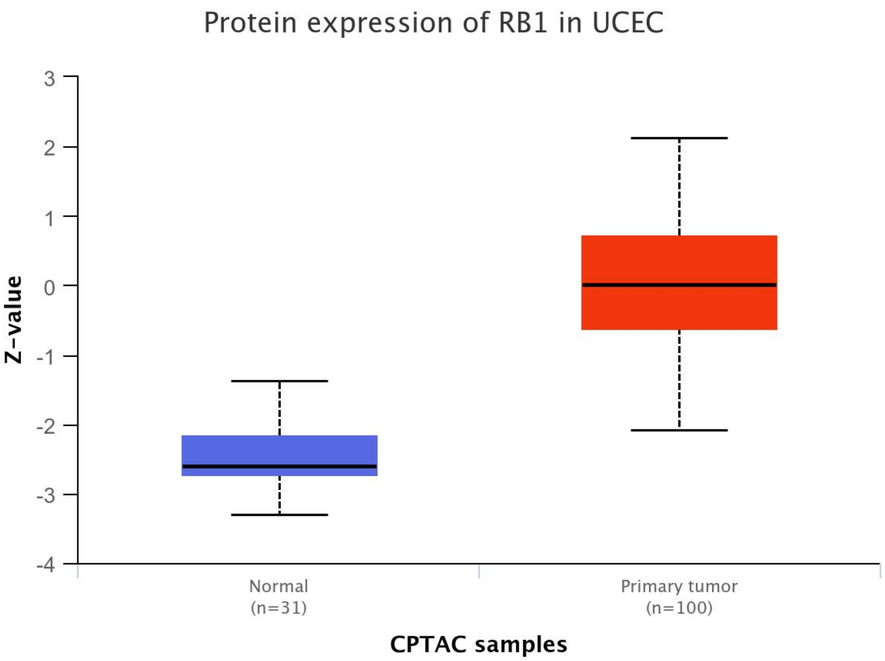

Protein expression of RB1 in UCEC
Z-value
3
2
1
0
-1
-2
-3
-4
Normal
(n=31)
Primary tumor
(n=100)
CPTAC samples

# S4.

A

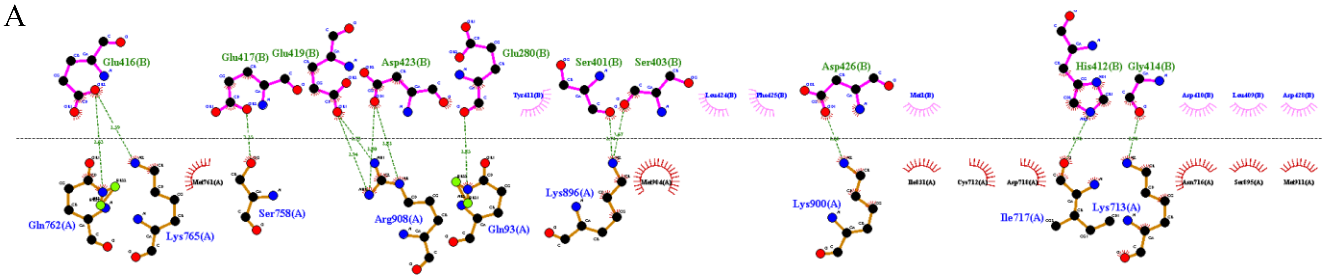


WTRB(E2F1)

B

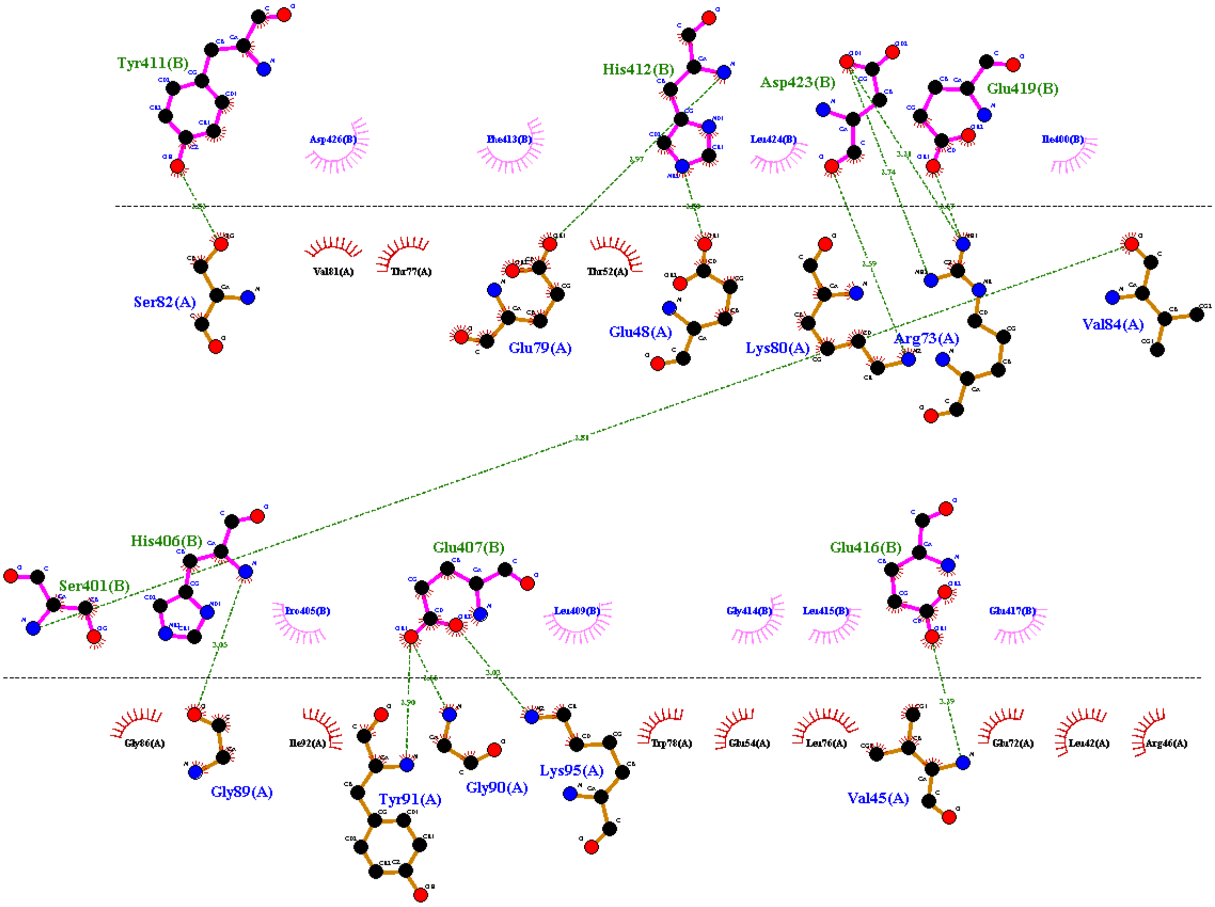


E137STOP(E2F1)

C

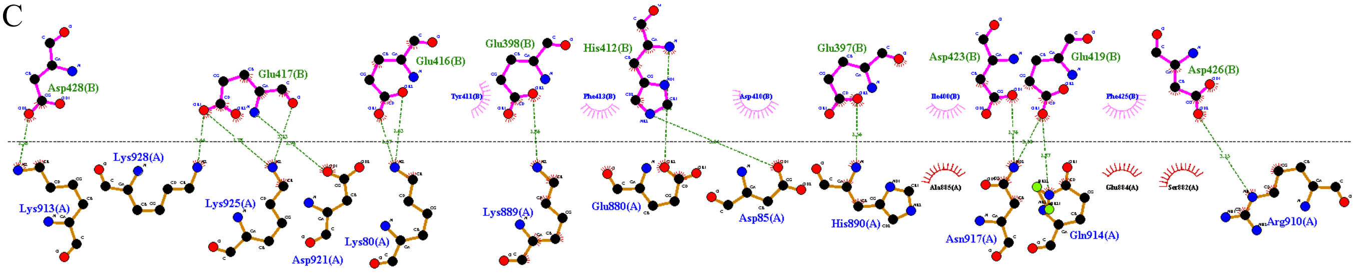


R272I(E2F1)

D

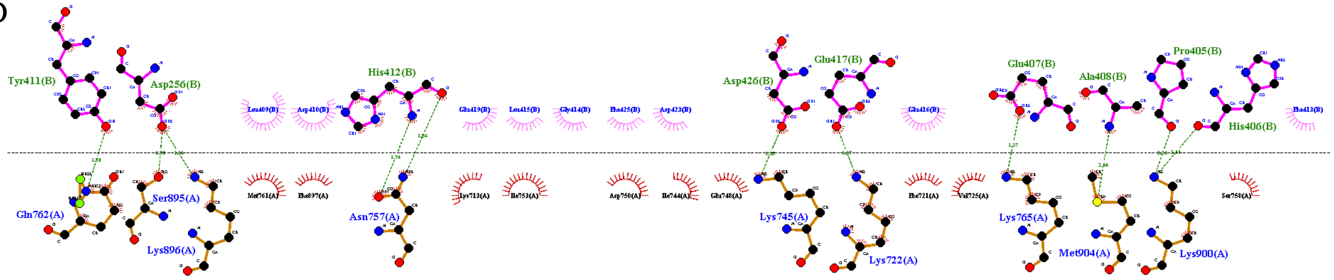


R741C(E2F1)

E

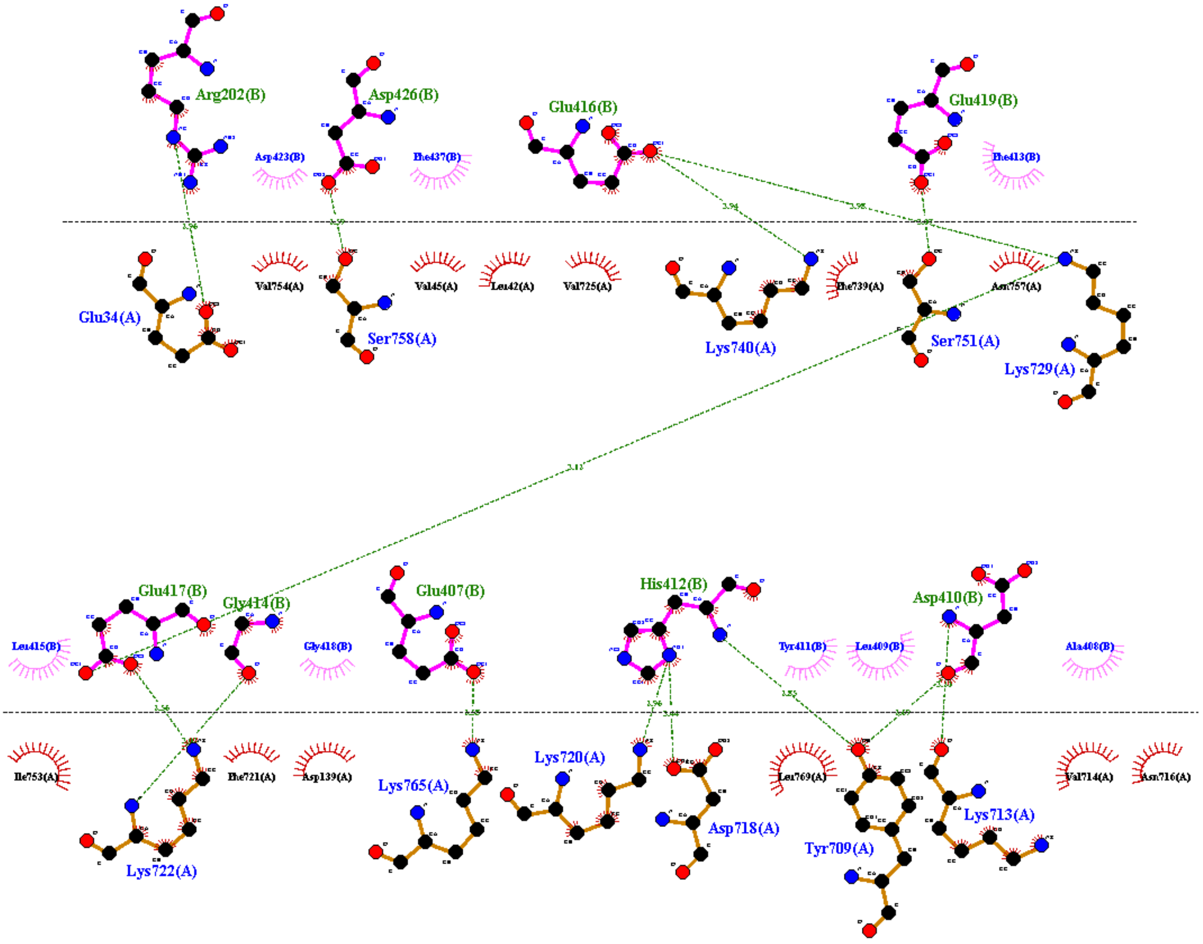


R876C(E2F1)

S5.

A

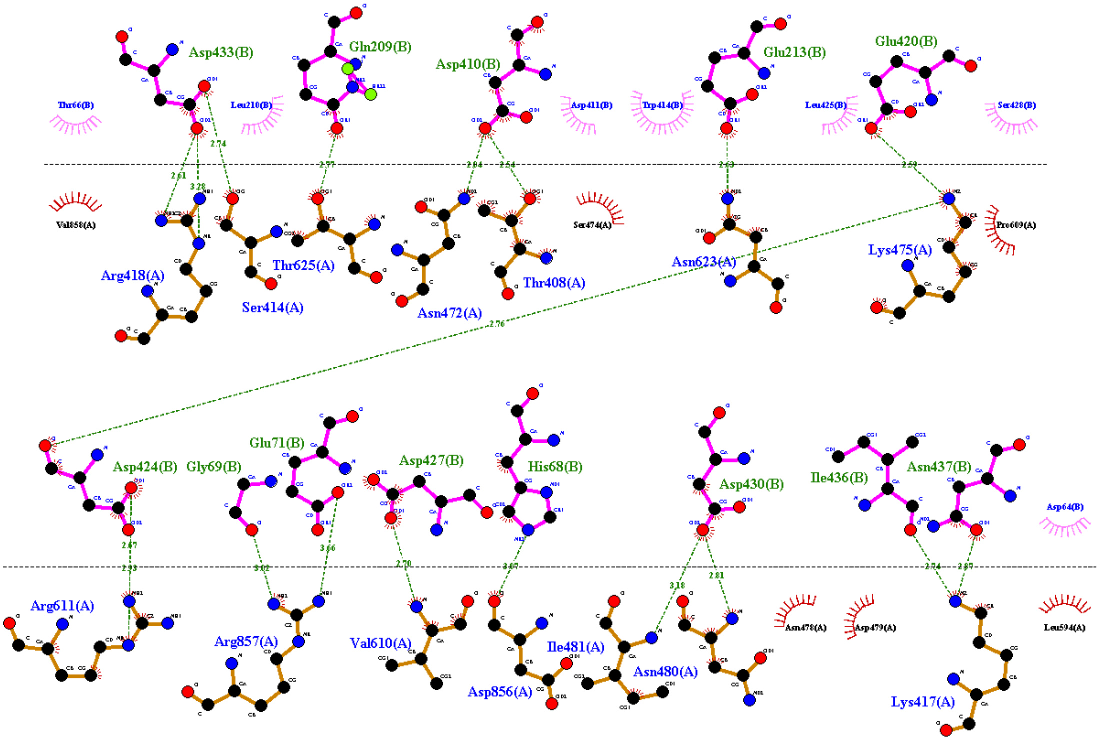


WTRB(E2F2)

B

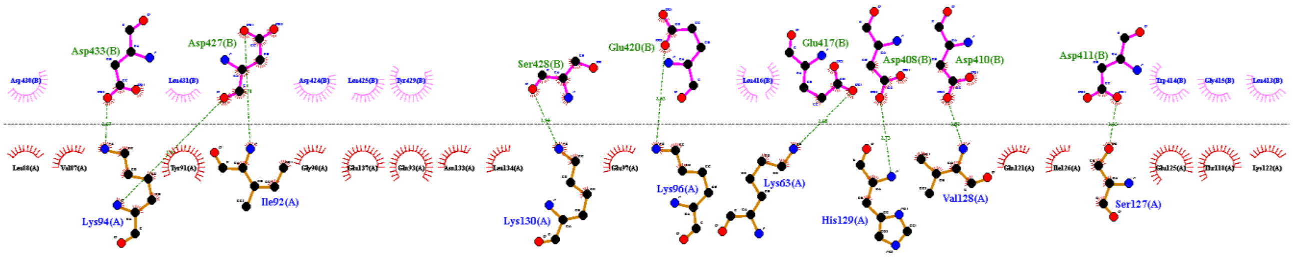


E137STOP(E2F2)

C

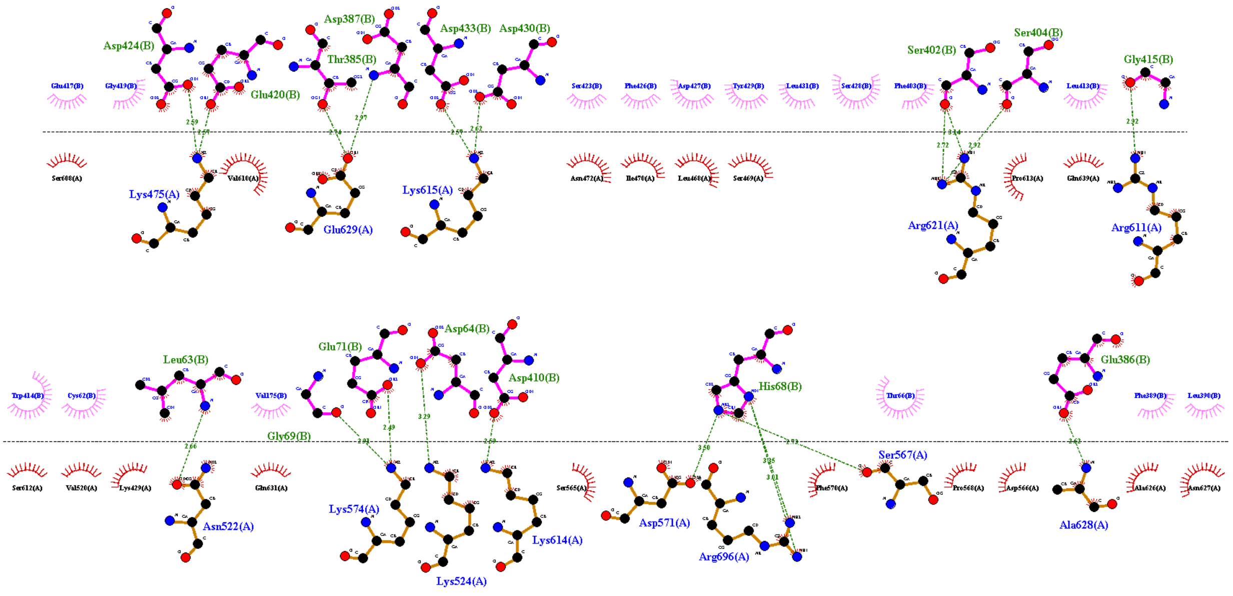


R272I(E2F2)

D

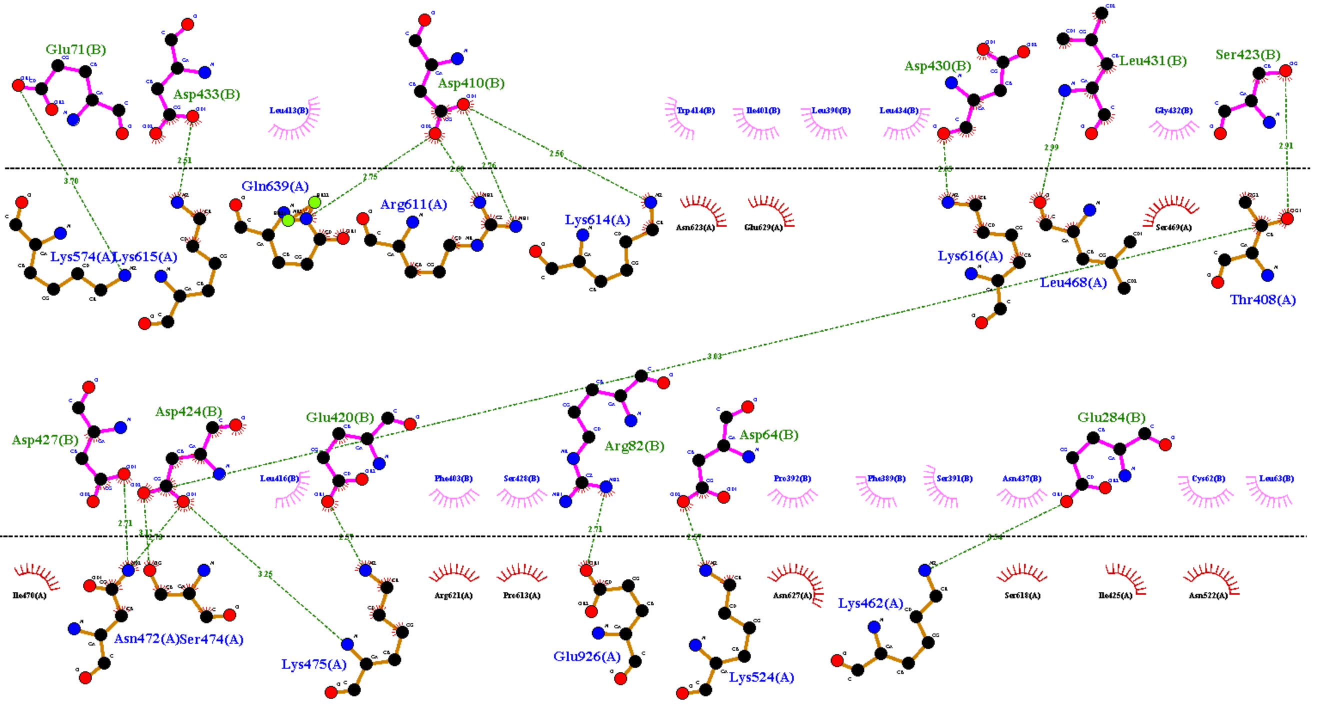


R741C(E2F2)

E

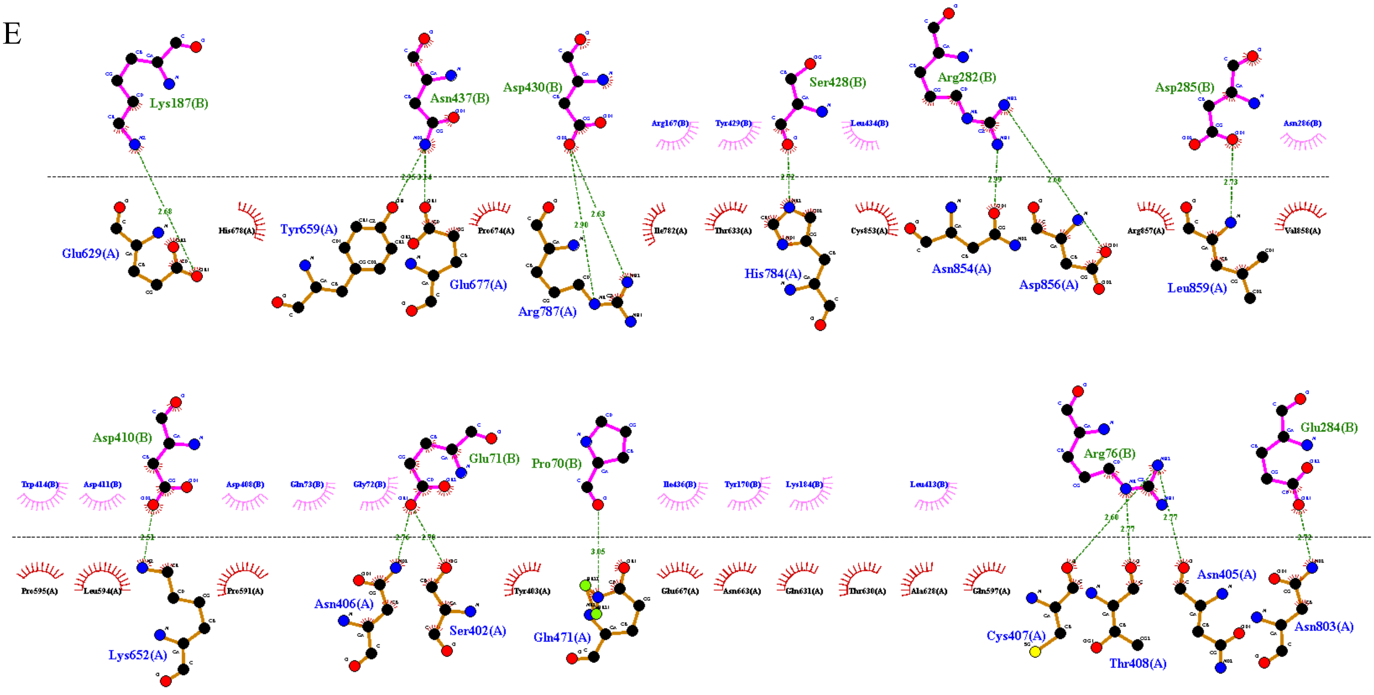


R876C(E2F2)

S6.

A

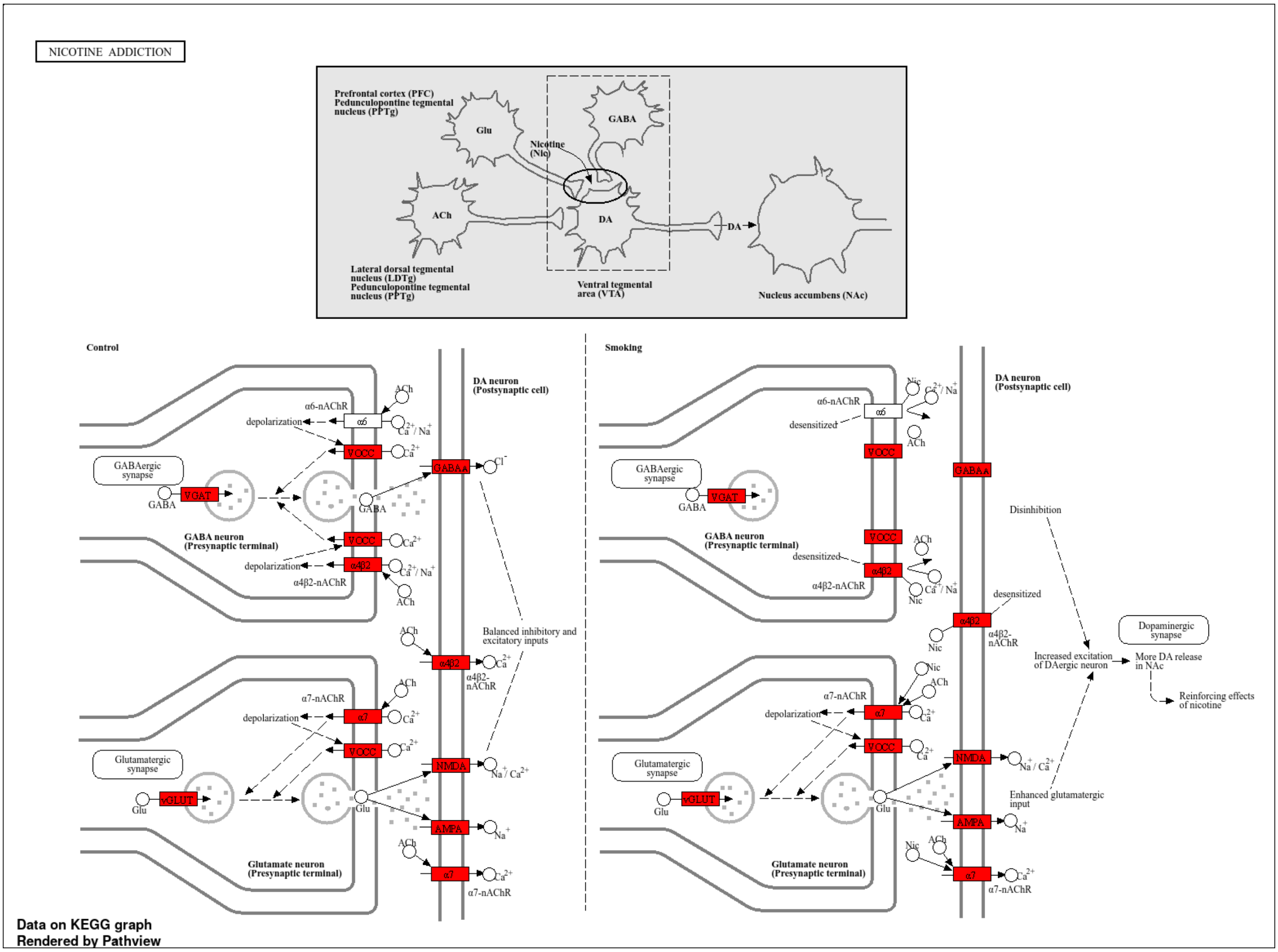
NICOTINE ADDICTION
Prefrontal cortex (PFC)
Pedunculopontine tegmental nucleus (PPTg)
Glu
GABA
Nicotine (Nic)
ACh
DA
DA
Lateral dorsal tegmental nucleus (LDTg)
Pedunculopontine tegmental nucleus (PPTg)
Ventral tegmental area (VTA)
Nucleus accumbens (NAc)
Control
Smoking
DA neuron (Postsynaptic cell)
GABAergic synapse
GABA neuron (Presynaptic terminal)
Glutamatergic synapse
Glutamate neuron (Presynaptic terminal)
α6-nAChR
α4β2-nAChR
α7-nAChR
depolarization
desensitized
VGAT
vGLUT
VOCC
GABAA
NMDA
AMPA
Balanced inhibitory and excitatory inputs
Disinhibition
Increased excitation of DAergic neuron
Enhanced glutamatergic input
Dopaminergic synapse
More DA release in NAc
Reinforcing effects of nicotine
Data on KEGG graph
Rendered by Pathview

B

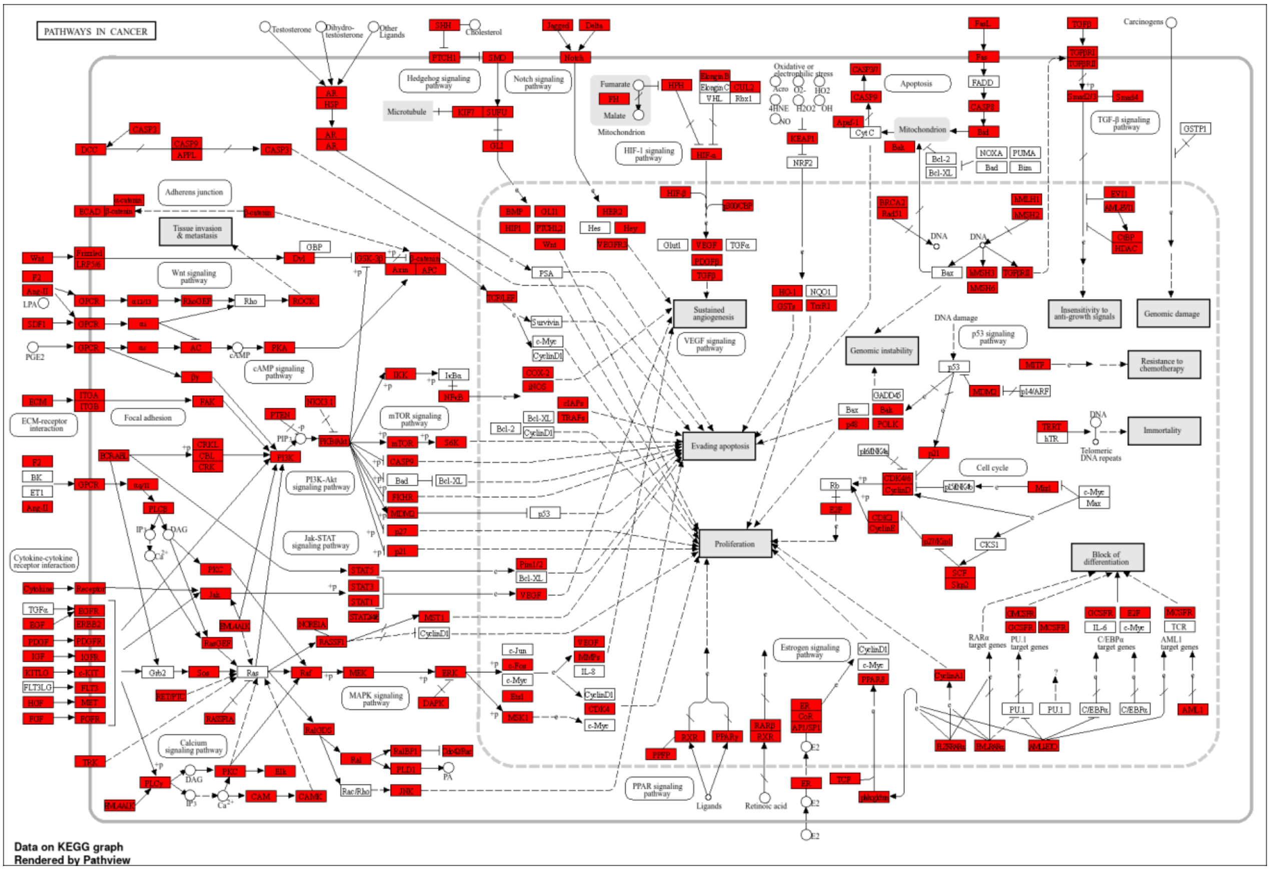
PATHWAYS IN CANCER
Data on KEGG graph
Rendered by Pathview

C

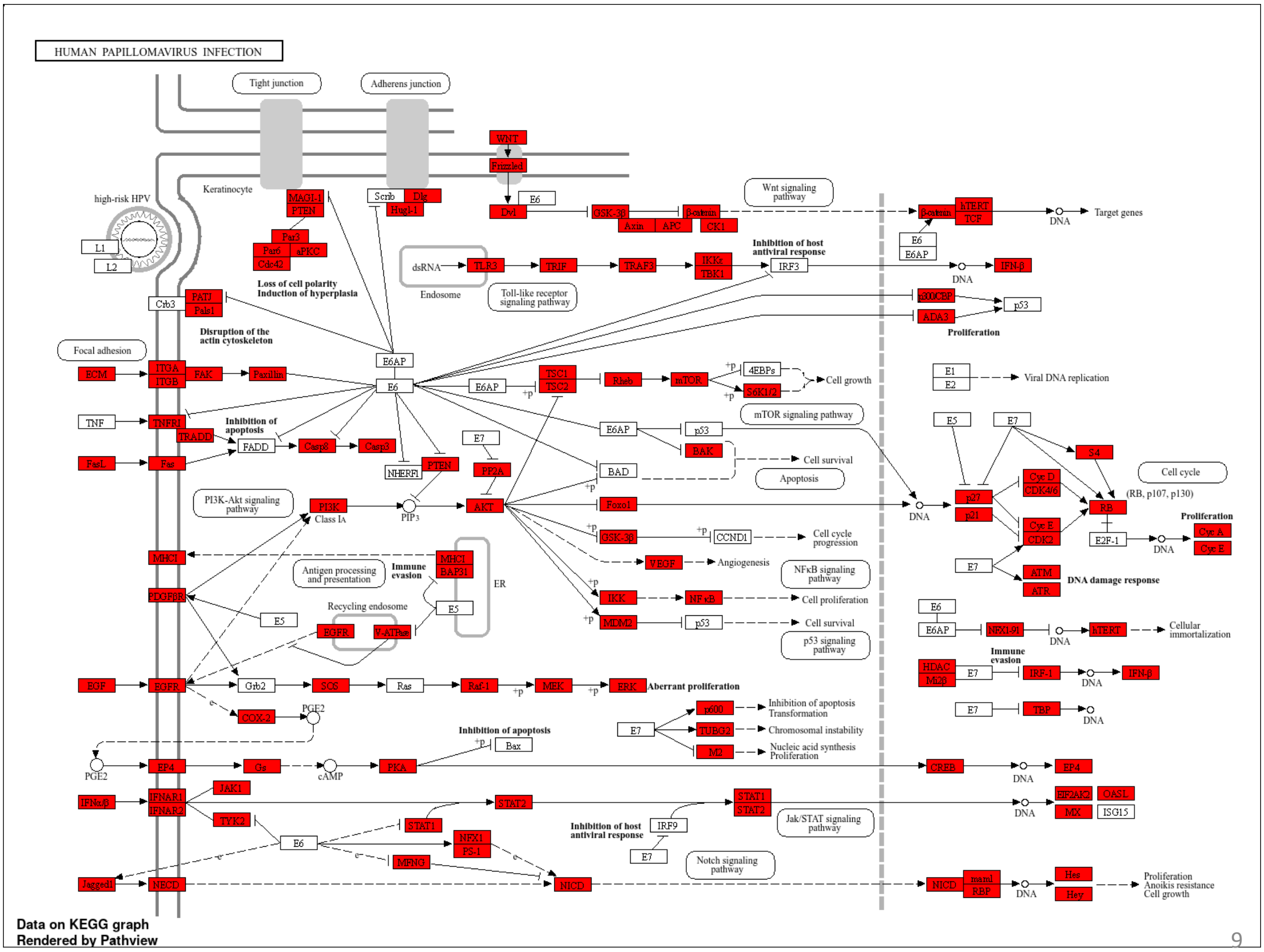
HUMAN PAPILLOMAVIRUS INFECTION
Tight junction
Adherens junction
high-risk HPV
Keratinocyte
Loss of cell polarity
Induction of hyperplasia
Disruption of the actin cytoskeleton
Inhibition of host antiviral response
Proliferation
Inhibition of apoptosis
Immune evasion
Aberrant proliferation
DNA damage response
Data on KEGG graph
Rendered by Pathview

S7.

A

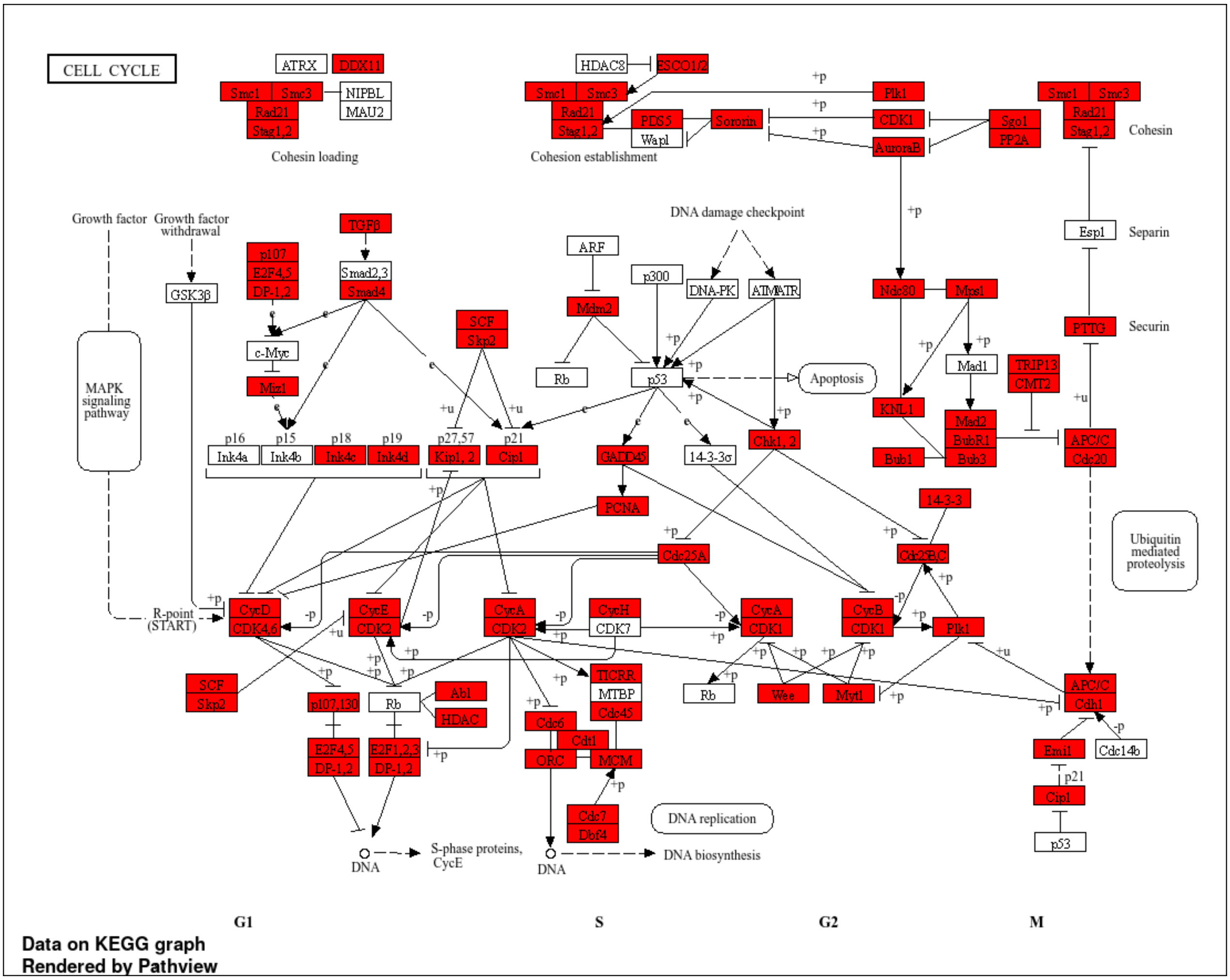
CELL CYCLE
ATRX
DDX11
Smc1
Smc3
Rad21
Stag1,2
NIPBL
MAU2
Cohesin loading
HDAC8
ESCO1/2
PDS5
Wapl
Sororin
Cohesion establishment
Plk1
CDK1
AuroraB
Sgo1
PP2A
Cohesin
Esp1
Separin
PTTG
Securin
TRIP13
CMT2
APC/C
Cdc20
Growth factor
Growth factor withdrawal
GSK3β
TGFβ
p107
E2F4,5
DP-1,2
Smad2,3
Smad4
c-Myc
Miz1
MAPK signaling pathway
SCF
Skp2
ARF
Mdm2
Rb
p300
DNA damage checkpoint
DNA-PK
ATM/ATR
p53
Apoptosis
Ndc80
Mps1
Mad1
KNL1
Mad2
BubR1
Bub3
Bub1
p16
p15
p18
p19
p27,57
p21
Ink4a
Ink4b
Ink4c
Ink4d
Kip1, 2
Cip1
GADD45
14-3-3σ
Chk1, 2
14-3-3
PCNA
Cdc25A
Cdc25B,C
Ubiquitin mediated proteolysis
R-point (START)
CycD
CDK4,6
CycE
CDK2
CycA
CycH
CDK7
CDK1
CycB
Plk1
Rb
Wee
Myt1
APC/C
Cdh1
Abl
HDAC
p107,130
E2F4,5
DP-1,2
E2F1,2,3
TICRR
MTBP
Cdc45
Cdc6
Cdt1
ORC
MCM
Cdc7
Dbf4
DNA replication
Emi1
Cdc14b
Cip1
p53
DNA
S-phase proteins, CycE
DNA biosynthesis
G1
S
G2
M
Data on KEGG graph
Rendered by Pathview

B

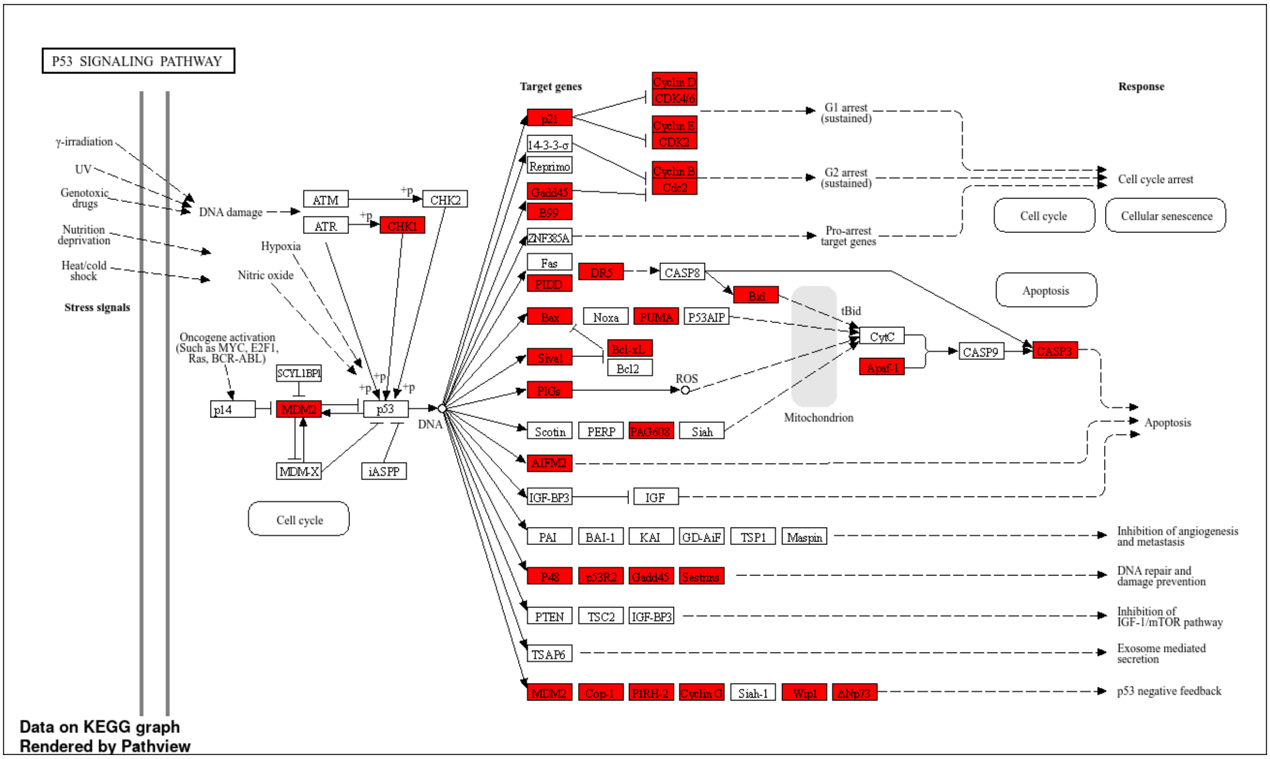
P53 SIGNALING PATHWAY
γ-irradiation
UV
Genotoxic drugs
Nutrition deprivation
Heat/cold shock
Stress signals
DNA damage
ATM
ATR
CHK2
CHK1
Hypoxia
Nitric oxide
Oncogene activation (Such as MYC, E2F1, Ras, BCR-ABL)
p14
SCYL1BP1
MDM2
MDM-X
p53
iASPP
DNA
Cell cycle
Target genes
p21
14-3-3-σ
Reprimo
Gadd45
B99
ZNF385A
Fas
PIDD
DR5
CASP8
Bid
tBid
Bax
Noxa
PUMA
P53AIP
Siva1
Bcl-xL
Bcl2
PIGs
ROS
CytC
Apaf-1
CASP9
CASP3
Mitochondrion
Scotin
PERP
PAG608
Siah
AIFM2
IGF-BP3
IGF
PAI
BAI-1
KAI
GD-AiF
TSP1
Maspin
P48
p53R2
Gadd45
Sestrins
PTEN
TSC2
IGF-BP3
TSAP6
MDM2
Cop-1
PIRH-2
Cyclin G
Siah-1
Wip1
ΔNp73
Cyclin D
CDK4/6
Cyclin E
CDK2
Cyclin B
Cdc2
G1 arrest (sustained)
G2 arrest (sustained)
Pro-arrest target genes
Response
Cell cycle arrest
Cell cycle
Cellular senescence
Apoptosis
Inhibition of angiogenesis and metastasis
DNA repair and damage prevention
Inhibition of IGF-1/mTOR pathway
Exosome mediated secretion
p53 negative feedback
Data on KEGG graph
Rendered by Pathview

C

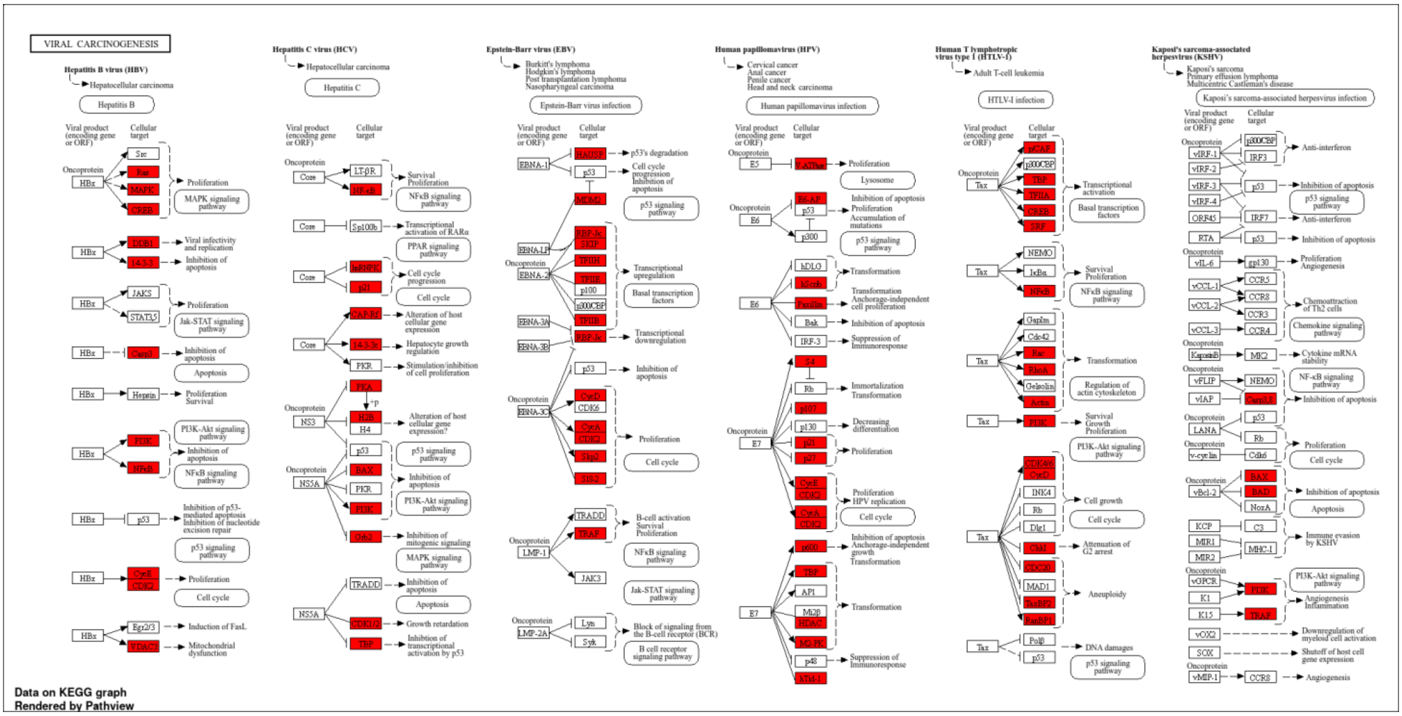
VIRAL CARCINOGENESIS
Hepatitis B virus (HBV)
Hepatitis C virus (HCV)
Epstein-Barr virus (EBV)
Human papillomavirus (HPV)
Human T lymphotropic virus type 1 (HTLV-1)
Kaposi's sarcoma-associated herpesvirus (KSHV)
Data on KEGG graph
Rendered by Pathview

S8.

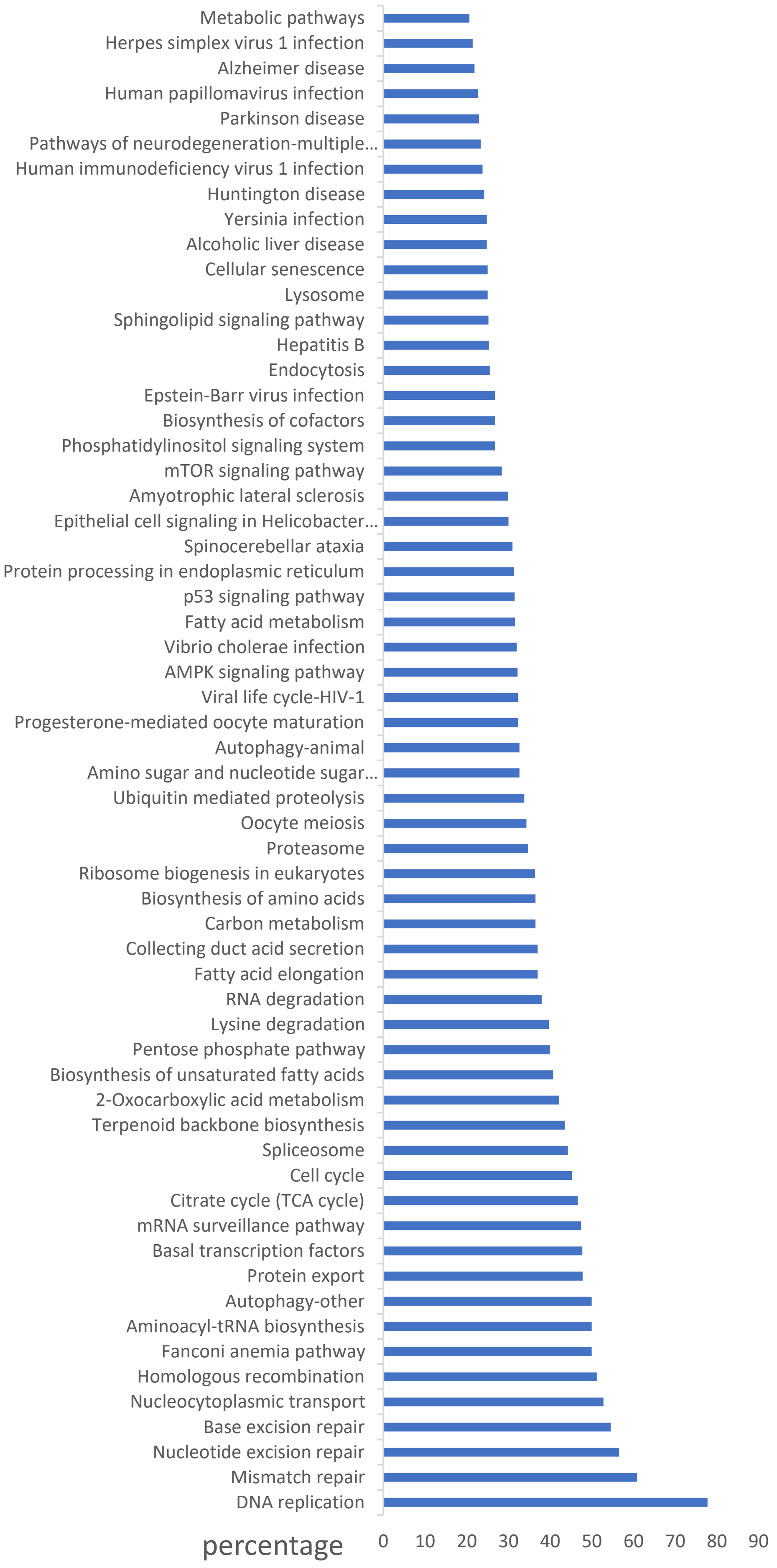

Metabolic pathways
Herpes simplex virus 1 infection
Alzheimer disease
Human papillomavirus infection
Parkinson disease
Pathways of neurodegeneration-multiple…
Human immunodeficiency virus 1 infection
Huntington disease
Yersinia infection
Alcoholic liver disease
Cellular senescence
Lysosome
Sphingolipid signaling pathway
Hepatitis B
Endocytosis
Epstein-Barr virus infection
Biosynthesis of cofactors
Phosphatidylinositol signaling system
mTOR signaling pathway
Amyotrophic lateral sclerosis
Epithelial cell signaling in Helicobacter…
Spinocerebellar ataxia
Protein processing in endoplasmic reticulum
p53 signaling pathway
Fatty acid metabolism
Vibrio cholerae infection
AMPK signaling pathway
Viral life cycle-HIV-1
Progesterone-mediated oocyte maturation
Autophagy-animal
Amino sugar and nucleotide sugar…
Ubiquitin mediated proteolysis
Oocyte meiosis
Proteasome
Ribosome biogenesis in eukaryotes
Biosynthesis of amino acids
Carbon metabolism
Collecting duct acid secretion
Fatty acid elongation
RNA degradation
Lysine degradation
Pentose phosphate pathway
Biosynthesis of unsaturated fatty acids
2-Oxocarboxylic acid metabolism
Terpenoid backbone biosynthesis
Spliceosome
Cell cycle
Citrate cycle (TCA cycle)
mRNA surveillance pathway
Basal transcription factors
Protein export
Autophagy-other
Aminoacyl-tRNA biosynthesis
Fanconi anemia pathway
Homologous recombination
Nucleocytoplasmic transport
Base excision repair
Nucleotide excision repair
Mismatch repair
DNA replication
percentage
0
10
20
30
40
50
60
70
80
90

Table 1

A.

| | WT | E137_Stop | R272I | R741C | R876C |
|---|---|---|---|---|---|
| HADDOCK score (kcal/mol) | -102.0 +/- 10.6 | -40.8 +/- 11.6 | -89.1 +/- 1.7 | -124.1 +/- 6.6 | -107.2 +/- 9.0 |
| RMSD from the overall lowest-energy structure (Å) | 2.2 +/- 1.4 | 19.6 +/- 1.2 | 41.1 +/- 0.2 | 10.0 +/- 0.3 | 41.8 +/- 1.2 |
| Van der Waals energy (kJ/mol) | -34.9 +/- 9.4 | -60.6 +/- 6.9 | -35.8 +/- 9.7 | -40.9 +/- 10.8 | -46.8 +/- 2.1 |
| Electrostatic energy (kJ/mol) | -407.9 +/- 60.0 | -332.6 +/- 16.4 | -412.0 +/- 54.0 | -511.0 +/- 66.0 | -346.1 +/- 54.2 |
| Desolvation energy (kJ/mol) | 8.9 +/- 4.7 | -1.1 +/- 2.4 | 15.4 +/- 5.3 | 17.9 +/- 4.4 | 7.8 +/- 6.1 |
| Restraints violation energy (kJ/mol) | 55.5 +/- 38.0 | 874.5 +/- 107.7 | 136.7 +/- 41.3 | 11.9 +/- 14.4 | 10.8 +/- 7.4 |
| Buried Surface Area (Å²) | 2024.0 +/- 183.0 | 1991.7 +/- 124.3 | 1806.6 +/- 114.5 | 2205.8 +/- 91.3 | 1966.6 +/- 116.9 |
| Z-Score | -1.9 | -1.8 | -1.4 | -1.5 | -1.9 |

B.

| | WT | E137_Stop | R272I | R741C | R876C |
|---|---|---|---|---|---|
| HADDOCK score (kcal/mol) | -98.4 +/- 21.8 | -16.9 +/- 18.6 | -138.6 +/- 5.4 | -103.4 +/- 30.4 | -135.5 +/- 7.5 |
| RMSD from the overall lowest-energy structure (Å) | 22.7 +/- 0.6 | 2.5 +/- 1.5 | 1.8 +/- 1.1 | 1.6 +/- 1.0 | 1.7 +/- 1.2 |
| Van der Waals energy (kJ/mol) | -42.6 +/- 3.8 | -41.2 +/- 11.9 | -49.1 +/- 9.6 | -37.6 +/- 4.9 | -81.5 +/- 7.2 |
| Electrostatic energy (kJ/mol) | -475.1 +/- 84.6 | -375.9 +/- 41.2 | -747.3 +/- 69.4 | -631.7 +/- 121.0 | -518.4 +/- 84.6 |
| Desolvation energy (kJ/mol) | 19.7 +/- 2.7 | 9.6 +/- 1.5 | 34.0 +/- 3.1 | 32.0 +/- 2.7 | 17.8 +/- 6.4 |
| Restraints violation energy (kJ/mol) | 195.8 +/- 87.9 | 899.2 +/- 87.3 | 260.1 +/- 43.3 | 285.9 +/- 68.7 | 318.6 +/- 75.2 |
| Buried Surface Area (Å²) | 2362.5 +/- 90.0 | 1629.4 +/- 145.8 | 2978.0 +/- 306.9 | 2576.3 +/- 249.8 | 3462.8 +/- 187.0 |
| Z-Score | -2.5 | -2.1 | -2.3 | -1.7 | -1.9 |
